# Multipotent radial glia sequentially deploy fate-restricted intermediate progenitors to diversify cortical interneurons

**DOI:** 10.1101/735019

**Authors:** Sean Michael Kelly, Ricardo Raudales, Monika Moissidis, Z. Josh Huang

## Abstract

GABAergic interneurons deploy numerous inhibitory mechanisms that regulate cortical circuit operation, but the developmental programs that generate diverse interneuron types are not yet well understood. We carried out a comprehensive genetic fate mapping of the radial glial progenitors (RGs) and intermediate progenitors (IP) in the medial ganglionic eminence (MGE) in mice. We reveal that *Nkx2.1^+^* RGs are multipotent and mediate two consecutive waves of neurogenesis, each sequentially generating different sets of interneuron types that laminate the neocortex in an inside-out-inside order. The first wave begins early and is restricted to the caudal MGE, has limited neurogenic window and capacity, and involves mostly apical IPs. The second wave begins later throughout the MGE, has an extended time course and large neurogenic capacity, and features a large set of fate-restricted basal IPs that amplify and diversify interneurons. Chandelier cells are generated toward the end of each wave and laminate in an outside-in order. Therefore, separate pools of multipotent RGs deploy temporal cohorts of IPs to sequentially generate diverse interneuron types.

## INTRODUCTION

In the mammalian neocortex, whereas glutamatergic pyramidal neurons (PNs) mediate myriad intra-cortical processing streams and output channels, GABAergic interneurons regulate the delicate balance and dynamic operations of functional neural ensembles (Harris and Shepherd 2015; Huang 2014). Although a minority compared to PNs, GABAergic neurons consist of diverse types with characteristic anatomical features, physiological properties, and gene expression profiles (Tremblay et al. 2016; Lim et al. 2018; Yuste et al. 2020; De Marco García and Fishell 2025; Huang and Paul 2019; van Velthoven et al. 2025; Lee et al. 2023; Gouwens et al. 2020). The diversity of GABAergic neurons is likely necessary for their functional versatility in shaping the spatiotemporal dynamics of circuit operations underlying cortical processing and information flow (Roux and Buzsáki 2015; Fishell and Kepecs 2020). However, the developmental programs that reliably and systematically generate such diverse interneuron types remain not well understood.

All cortical GABAergic neurons derive from the embryonic ventral telencephalon, or subpallium, which comprises several subdivisions such as the medial and caudal ganglionic eminence (MGE and CGE, respectively) and the preoptic area (POA) (Hu et al. 2017; Kepecs and Fishell 2014; Lim et al. 2018; Sultan and Shi 2018; Bandler and Mayer 2023). Among these, MGE and POA are marked by the homeodomain protein NKX2.1 and give rise to major subclasses such as parvalbumin (PV) and somatostatin (SST) expressing interneurons (Sussel et al. 1999; Hu et al. 2017). In addition, there is evidence for finer regionalization along the dorsal-ventral axis of MGE-POA that may bias the production of PV versus SST neurons (Flames et al. 2007; Hu et al. 2017; Long et al. 2009). Furthermore, developmental timing (e.g. time of cell birth) also influences interneuron specification and laminar position (Miyoshi et al. 2007; Bright et al. 2025). At the cellular level, the neuroepithelium within MGE/POA consists of diverse types of neural progenitors, including regenerative radial glial cells (RGs), which give rise to apical as well as basal intermediate progenitors (aIP and bIPs, respectively) (Brown et al. 2011; Ciceri et al. 2013; Harwell et al. 2015; Mayer et al. 2015; Turrero García and Harwell 2017). More recent studies suggest that MGE/POA RGs are multi-potent and sequentially generate multiple types of cortical interneurons (Sultan et al. 2018; Sultan and Shi 2018).

Despite this significant progress, we are yet to achieve a cellular resolution lineage framework that incorporates the spatial and temporal aspects of MGE neurogenesis while linking progenitor types and their lineage progression to diverse interneuron types. For example, it remains unresolved whether an initial “founding pool” of RGs has the potential to generate all MGE-derived GABAergic neuron types, or whether separate RG pools, each multi-potent to varying degrees, give rise to distinct subsets and/or ratios of interneurons (Hu et al. 2017). Moreover, the role of IPs is largely unexplored, thus the relationship of RGs to different IP types and to interneuron types is not well understood.

Here, we combined multiple intersectional recombinase driver lines to systematically fate-map MGE/POA RGs and IPs starting from the onset and throughout the course of neurogenesis. We further birthdated and tracked their respective interneuron progeny, resolving the laminar and morphological types in the mature cortex. Surprisingly, we found that the MGE comprises two separate RG pools with distinct spatiotemporal progression patterns, IP types, neurogenic capacities and outputs. Together, these pools mediate two waves of neurogenesis, each sequentially generating different sets of interneuron types that laminate the neocortex in an inside-out-inside sequence. Whereas a small and early-activated RG pool (at ∼embryonic day E10) resides in the caudal MGE and involves mostly *Ascl1^+^*aIPs with limited interneuron output, a large RG pool, initiated later at ∼E12, populates the entire rostral-caudal extent of MGE and features temporal cohorts of fate-restricted *Ascl1^+^*and *Dlx1^+^* bIPs that amplify and diversify interneuron production. In particular, we find that chandelier cells (ChCs) are generated toward the end of each wave and laminate the cortex in an outside-in order. These results establish a cellular resolution lineage framework in the MGE whereby multipotent RGs deploy distinct temporal subsets of fate-restricted IPs to sequentially amplify and diversify interneuron types.

## RESULTS

### The MGE comprises two separate RG pools with distinct spatiotemporal characteristics and neurogenic capacity

Upon initial formation at ∼E9.5, MGE is delineated by NKX2.1 expression within a small region of the subpallial ventricular zone (VZ) (Sussel et al. 1999), which consists of a thin layer of neuroepithelial cells (NEs). NEs then proliferate and differentiate into RGs, which mediate neurogenesis either directly or indirectly through IPs (Brown et al. 2011). MGE neurogenesis begins soon after E9.5 (Miyoshi et al. 2007; Sussel et al. 1999), peaks between E12-15, and continues until ∼E17.5 (Sultan et al. 2018). During this period, the initially inconspicuous VZ expands dramatically along the rostral-caudal, dorsal-ventral, and apical-basal axes. In parallel, the MGE differentiates into multiple subregions delineated by combinatorial transcription factor expression (Flames et al. 2007; Long et al. 2009), giving rise to numerous dorsal-ventral subdomains and an increasingly thick subventricular zone (SVZ) layer deep to the VZ (Petryniak et al. 2007). Using a retrovirus-based fate mapping approach, a previous study demonstrated that, as early as E12.5, MGE RGs are multi-potent to generate diverse cortical interneurons across layers 2 to 6 (Sultan et al. 2018). However, the properties of the initial founding population of RGs and their relationship to subsequent RGs have not been explored, leaving a major gap in our understanding of the MGE lineage. Here, we used the *Nkx2.1-CreER* knockin driver line (Taniguchi et al. 2011) to fate map MGE progenitors from the very onset and throughout the course of neurogenesis. Through pulse-chase labeling of these progenitors across multiple embryonic times up until adulthood, progenitor characteristics were tracked including lineage progression and interneuron output.

Tamoxifen (TM) induction at E10 followed by 8-hour pulse-chase in *Nkx2.1-CreER;Ai14* mice labeled the earliest set of RGs throughout the nascent MGE at this stage (Figure 1A-E). These RGs, which were characterized by end-feet attached to the ventricular surface and a notable basal fiber (Figure 1C, D; Figure S1A), are hereafter designated as RG^E^s. In addition, putative apical IPs (aIPs; with ventricular end-feet but no basal fiber) were also labeled (Figure 1F, S1B,E). Whereas RGs generally expressed low or undetectable levels of the proneural factor ASCL1, aIPs expressed medium to high levels of ASCL1 (Figure S1B, E, G), corresponding with their subsequently more active participation in neurogenesis. Interestingly, the *Nkx2.1-CreER* driver appeared to preferentially label progenitors with higher levels of NKX2.1 immunoreactivity (Figure S1A), while the *Ascl1-CreER* driver labeled progenitors with both high and low levels of NKX2.1 immunoreactivity (Figure S1D). Both drivers labeled dividing progenitors but not postmitotic neurons (Figure S1C,F).

**Figure 1.**
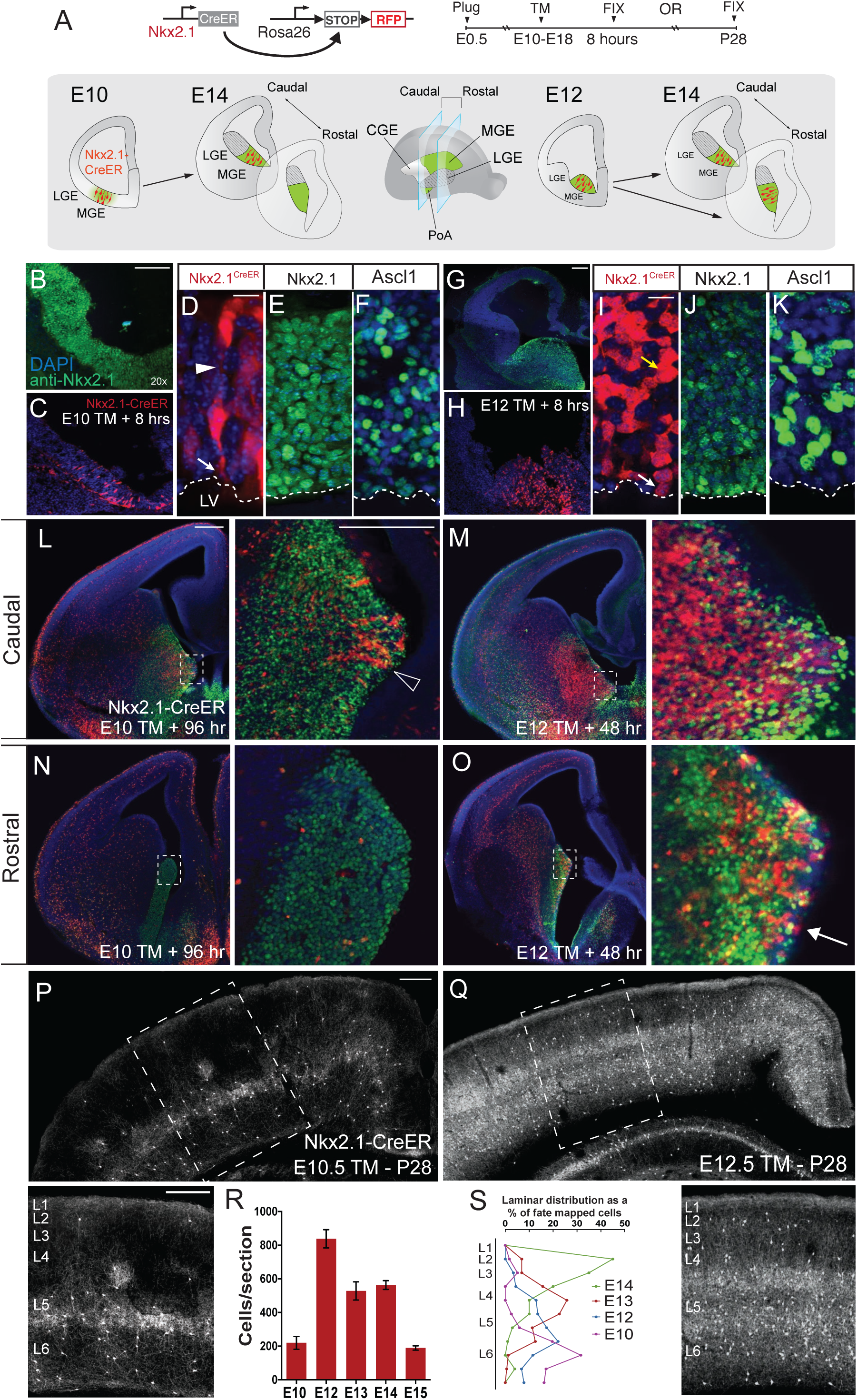
The MGE comprises two radial glial progenitor pools with distinct spatiotemporal characteristics and neurogenic capacity. (A) Strategy for genetic fate mapping of MGE RGs. Lower schematics depict the spatial restriction of the early E10 progenitors and their descendants which remain in the caudal domain of MGE even by E14 (left); in contrast, the later E12 progenitors and their descendants spread broadly to both rostral and caudal MGE (right). The levels of caudal and rostral coronal planes are indicated (middle). (B) E10 MGE marked by anti-NKX2.1 immunofluorescence (green). (C) E10 RGs sparsely labeled by 8-hour TM pulse-chase using the *Nkx2.1-CreER;Ai14* mice. (D-F) Magnified view of *Nkx2.1-CreER*-labeled E10 RGs showing the endfoot (D, arrow) at the lateral ventricle (LV) and radial process (arrowhead), as well as the apical to basal distribution of NKX2.1 (E) and ASCL1 immunoreactivity (F) within the early MGE ventricular zone. (G) E12 MGE marked by anti-NKX2.1 immunofluorescence, showing the bulging shape and increased thickness of the SVZ. (H) E12 MGE progenitors labeled by 8 hour TM pulse-chase. (I-K) High magnification view of E12 MGE showing the presence of RG with endfoot (I, white arrow) at the lateral ventricle (LV) and putative IPs in the SVZ (yellow arrow). The apical to basal distribution of NKX2.1 (J) and ASCL1 (K) immunoreactivity extending from the LV is shown in green. Note the apical to basal gradient of Nkx2.1 immunoreactivity, while ASCL1 immunoreactivity is more sporadic in its distribution in the SVZ. (L, N) Pulse-chase labeled E10 MGE progenitors and their descendants remain strictly confined to the caudal region after 96 hours (arrowhead indicates RG endfeet), without extending into the rostral domain of MGE (N). Left subpanels are coronal embryonic sections; dashed boxes represent the location of the high-magnification views in right subpanels. MGE is marked by NKX2.1 immunofluorescence (green). (M, O) Pulse-chase labeled E12 MGE progenitors and their descendants distribute broadly throughout both the caudal (M) and rostral (O) domain of the MGE (RG endfeet indicated by arrow). (P) Fate mapping of E10 MGE progenitors in *Nkx2.1-CreER;Ai14* mice shows that RG^E^s do not generate the full complement of interneurons that occupy all cortical layers, but instead produce subpopulations in L6, L5, and to a much more limited extent L2/3, with a conspicuous paucity of cells in L4. Representative images of somatosensory cortex at P28 are shown. Dashed box represents the location of the higher magnification subpanel below. (Q) In striking contrast, fate mapping of E12 RG^L^s shows that they generate a massive and broadly fated set of interneurons distributed throughout L2-6, including a rich assortment in L4. (R) Quantification of total cortical cell output in selected regions in mPFC, MC and SSC from *Nkx2.1-CreER* fate mapping between E10-15, quantified by number of cells per section with n = 6 sections from 3 separate brains for each time point. (S) Laminar distribution of fate-mapped interneurons in SSC from different embryonic days. Graph shows the percent distribution of cells quantified into 10 serial bins from pial surface to white matter. Scale bars: B-C: 100 um; D-F: 10 um; G-H: 200 um; I-K: 10 um L-O: 200 um in left panels, 100 um in right panels; P-Q: 100 um in top and bottom panels.

To follow the spatial and temporal progression of RG^E^s within the MGE, we performed 48-hour (i.e. to E12) and 96-hour (i.e. to E14) pulse-chase after E10 TM induction in *Nkx2.1-CreER;Ai14* mice. Labeled RGs and postmitotic neurons persisted in the VZ at these later times, indicating the regenerative properties of RG^E^s (Figure 1L). The descendants of E10 RG^E^s at E12 and E14 were detected only in the caudal region but not the rostral region of the substantially expanded MGE (Figure 1L, N), suggesting a distinct spatial restriction of the RG^E^ lineage within the MGE.

To reveal the neuronal output from these RG^E^s, we assayed the somatosensory cortex at P28 after E10 TM induction (Figure 1P, R). We found that RG^E^s did not give rise to interneurons across layers 2-6; instead, their progeny were detected mostly in infragranular layers, especially layer 5 (L5), and only sparsely in L2 and 3 (Figure 1P, S). Overall, a rather small number and subset of interneurons were consistently labeled, with a conspicuous absence of labeling in L4 and much of L2/3. This result was highly reproducible across experiments in more than 10 litters of animals independent of high or low TM dose. Interestingly, similar results on such a limited capacity and infragranular-biased output of early RGs were also observed by retrovirus-mediated fate mapping at E11.5 (Ciceri et al. 2013). Together, these results suggest that the earliest set of MGE RGs restricted their lineage progression in the caudal MGE and had quite limited neurogenic capacity.

In stark contrast, equivalent fate mapping experiments in *Nkx2.1-CreER;Ai14* mice at E12 yielded a very different outcome. In this case, 8-hour pulse-chase at E12 not only labeled a large set of RGs in the VZ, hereafter designated as RG^L^s (Figure 1G-J), but further captured a large population of basal IPs (bIPs) in the SVZ with higher levels of ASCL1 immunoreactivity (Figure 1I-K, Figure S1H). At 48-hour (i.e. to E14) after E12 TM induction, labeled RGs and postmitotic neurons persisted in the VZ, again indicating the self-renewing properties of RG^L^s (Figure 1M). Importantly however, the descendants of RG^L^s at E14 occupied the entire caudal-to-rostral axis of MGE, as determined by co-immunolabeling with anti-NKX2.1 antibodies (Figure 1M,O), in contrast to the caudally-restricted descendants of RG^E^s.

At P28, RG^L^-derived interneurons were consistently exceptionally abundant and settled across layers 2 to 6 (Figure 1Q-S), also in contrast to RG^E^-derived interneurons. Further fate mapping of *Nkx2.1^+^* progenitors with TM pulses on E13 and E14 gave rise to progressively greater numbers of supragranular interneurons (Figure 1R,S), which were likely subsets of those generated from E12 RG^L^s. These fate mapping results at E12, E13, E14 were consistent with those obtained using retrovirus-mediated fate mapping at the same embryonic times (Sultan et al. 2018). Together, these results suggest that the early MGE comprises two separate pools of RGs. Whereas RG^E^s are restricted to the caudal MGE and generate a limited number and subsets of interneurons, RG^L^s occupy the entire MGE and generate much more numerous and diverse interneurons across L2-L6.

### RG^E^s and RG^L^s are multipotent and generate interneurons that laminate the cortex in an inside-out-inside sequence

To probe the lineage relationship among interneurons derived from RG^E^s and RG^L^s, we combined fate mapping in *Nkx2.1-CreER;Ai14* mice with thymidine-analogue neuronal birth dating. Namely, E10 TM induction followed by E11 EdU injection and E13 BrdU injection labeled descendant neurons with RFP and those born at E11 and E13 with EdU and BrdU, respectively (Figure 2A; under RG^E^). This revealed that fate-mapped L6, but not L2/3 neurons were EdU^+^ (Figure 2B), while fate-mapped L2/3 and another set of L5/6 neurons were often BrdU^+^ (Figure 2D-F). We also detected a small number of EdU^+^/BrdU^+^ cells (Figure 2C), indicating that they derived from a single RG^E^ that divided on both E11 and E13.

**Figure 2.**
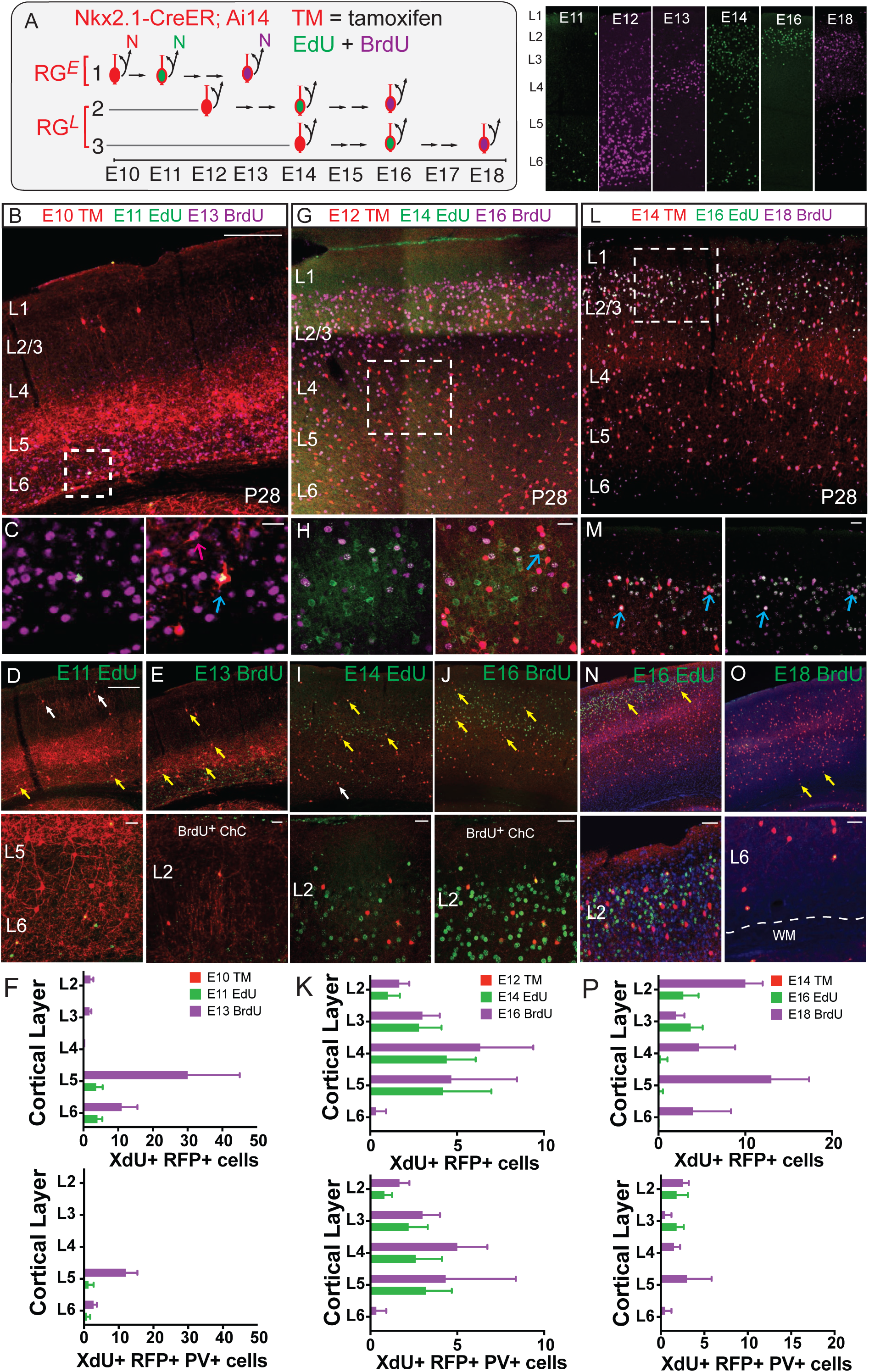
RG^E^s and RG^L^s are multipotent and generate interneurons that laminate cortex in an inside-out-inside sequence. (A) Experimental design for combined fate mapping and sequential birth dating of MGE interneurons. After TM induction of pregnant *Nkx2.1-CreER;Ai14* mice at E10, E12, or E14, sequential pulses of EdU and BrdU were given as indicated. Right panel: examples of the cortical distribution of EdU^+^ and BrdU^+^ cells at P28 following pulses on indicated embryonic days. (B-E) RG^E^s are multipotent and generate interneurons that laminate in an inside-out-inside sequence. E10 fate-mapped RFP^+^ neurons populated L6, L5 and L3. E11 EdU pulses co–labeled L6 (blue arrow in C; yellow arrows in D), but not L3 (white arrows in B, D) neurons. In contrast E13 BrdU pulses preferentially labeled L3 as well as additional L5 neurons (yellow arrows in E). An example of a L6 triple-labeled interneuron derived from a progenitor that sequentially divided to incorporate both EdU (green) on E11 and BrdU (magenta) on E13 is shown in (C), enlarged from the dashed box in (B). Examples of ChCs from RG^E^s are shown in bottom panel of (E). (F) Quantification of the laminar distribution of E10 fate-mapped and birth-dated total (upper graph) and PV^+^ (lower graph) interneurons. The graph shows number of cells per section. Note that of L6 neurons are born on E11, while proportionately more L5 neurons are born on E13, and L3 cells are exclusively co-labeled on E13 along with the deep layer cells, suggesting an inside-out-inside sequence. Non-ChC PV^+^ interneurons birth-dated with this cohort localize exclusively to cortical layers 5 and 6. (G-J) RG^L^s are multipotent and generate interneurons that laminate in an inside-out trend during mid-gestation. E12 fate-mapped interneurons populate each layer from L6 to L2. E14 EdU pulses co-labeled L3 and L4 cells (yellow arrows in I), but were never found to co-label L6 cells (white arrow in Ix), while E16 BrdU pulses co-labeled L2/3 as well as some L5 neurons (yellow arrows in J). Examples of triple-labeled cells in middle cortical layers are shown in (H, blue arrow). Examples of L2 ChCs at the L1/2 border produced by RG^L^s are shown in bottom panels of I, J. (K) Quantification of the laminar distribution of E12 fate-mapped and birth-dated total (upper graph) and PV^+^ (lower graph) interneurons. (L-O) During late phase neurogenesis, RG^L^s generate interneurons that laminate in an outside-in pattern. E14 fate-mapped neurons were distributed across layers 6 to 2. Whereas E16 EdU pulses co-labeled L2/3 cells almost exclusively (yellow arrows in N), E18 BrdU pulses co-labeled a significant number of L5/6 neurons in addition to L2/3 neurons (yellow arrows in O, lower panels). Examples of triple-labeled cells in L2/3 are shown in (M, blue arrows). (P) Laminar distribution of E14 fate-mapped and birth-dated total (upper) and PV^+^ (lower) interneurons. Note that at E18 a significant number of L5/6 neurons were generated in addition to L2/3 neurons. Scale bars: B, G, L: 300 um; C, H, M: 10 um; D, E, I, J, N, O: 300 um in top panels and 10 um in bottom panels

Further probing the subclass identity of these cells with immunohistochemistry, we found that more PV^+^ cells were born at E13 than at E11 (Figure 2F, Figure S2A,B). Also of note, we detected a small number of supragranular ChCs that were born selectively on E13 (Figure 2E, bottom subpanel). Together, these results suggest that RG^E^s are multipotent and sequentially generate a set of interneurons that are deployed to the cortex, first in deep layers and then in superficial layers as well as deep layers.

To further probe the temporal and laminar pattern of interneuron production from RG^L^s, we designed two additional EdU/BrdU labeling schemes (Figure 2A; under RG^L^). We found that of the massive number of E12 RG^L^s-derived and RFP-labeled cells across L6 to L2, middle to more supragranular cells were born at E14, while both supragranular and infragranular cells were born at E16 (Figure 2G-K). Interestingly, sequential E16 EdU and E18 BrdU labeling revealed that whereas E16-born cells mostly resided in supragranular layers, a significant fraction of E18-born cells occupied infragranular L5 and L6 (Figure 2L-P). Moreover, we detected a number of EdU^+^/BrdU^+^ cells (Figure 2M), indicating that they derived from the same RGs that divided at E14 and E16. In particular, PV^+^ cells were born throughout RG^L^ neurogenesis and laminated in an inside-out-inside pattern (Figure 2P, Figure S2C-F). Together these results suggest that RG^L^s are multipotent and sequentially generate a large set of interneurons that also laminate the cortex in an inside-out-inside (i.e. inside to outside and to inside again) pattern.

### IPs amplify laminar-restricted interneuron production in an inside-out-inside sequence

Previous studies demonstrate that MGE RGs undergo asymmetric divisions to self-renew and generate either a neuron or an IP while remaining attached to the ventricle surface (Brown et al. 2011; Turrero García and Harwell 2017). IPs then undergo symmetric divisions to generate pairs of neurons and comprise two major types: aIPs extend end-feet to the ventricle surface and reside in the VZ, whereas bIPs move to the subventricular zone (SVZ) before their neurogenic divisions (Pilz et al. 2013; Turrero García and Harwell 2017). The rapid expansion of SVZ from E12 onward suggests that the large majority of interneurons are likely generated through IPs. However, the precise role of IPs in the amplification and diversification of interneurons is not well understood.

While all subpallial IPs are thought to express *Ascl1*, many also express the homeodomain transcription factor *Dlx1* (Long et al. 2009; Wang et al. 2013). *Dlx1* may act downstream as well as in parallel to *Ascl1*, and IPs may contain *Ascl1^+^*, *Dlx1^+^*, and *Ascl1^+^*/*Dlx1^+^*subpopulations at different stages (Wang et al. 2013). We thus used the *Ascl1-CreER* (Sugahara et al. 2016) and *Dlx1-CreER (Taniguchi et al. 2011)* drivers to characterize IPs in the MGE across the course of neurogenesis (Figure 3A). At E10, 8-hour pulse-chase in *Ascl1-CreER;Ai14* mice labeled abundant aIPs in the VZ of MGE with characteristic end-feet attached to the ventricle and the absence of basal processes, distinct from RGs (Figure 3B). Importantly, equivalent experiments in *Dlx1-CreER;Ai14* mice labeled *only* postmitotic neurons in the mantle zone (MZ) but not IPs, indicating that early aIPs do not express *Dlx1*, which at this stage is confined to postmitotic neurons only (Figure 3C). In contrast, from E12 to E17 8-hour pulse-chase in *Ascl1-* and *Dlx1-CreER* drivers labeled almost exclusively bIPs in the SVZ (Figure 3D-G), hereafter designated as bIP*^Ascl1^* and bIP*^Dlx1^*, respectively. Initially, between E12-E13, we observed bIPs*^Ascl1^* and bIPs*^Dlx1^* to occupy the entire ventral-to-dorsal axis of the MGE and overlapped extensively, suggesting that most IPs during this period may express both genes (Figure 3D,E; (Wang et al. 2013)). Their distribution patterns then diverged from E14-E17, such that bIPs*^Dlx1^*and bIPs*^Ascl1^* were more concentrated in the dorsal and ventral MGE, respectively (Figure 3F,G). Together, these results suggest the progression of different spatial and temporal cohorts of IP types during MGE neurogenesis, from early aIPs *^Ascl1^* in the caudal region to broadly distributed bIPs*^Ascl1/Dlx1^* and to ventral-dorsal segregated bIPs*^Ascl1^* and bIPs*^Dlx1^*in the late MGE.

**Figure 3.**
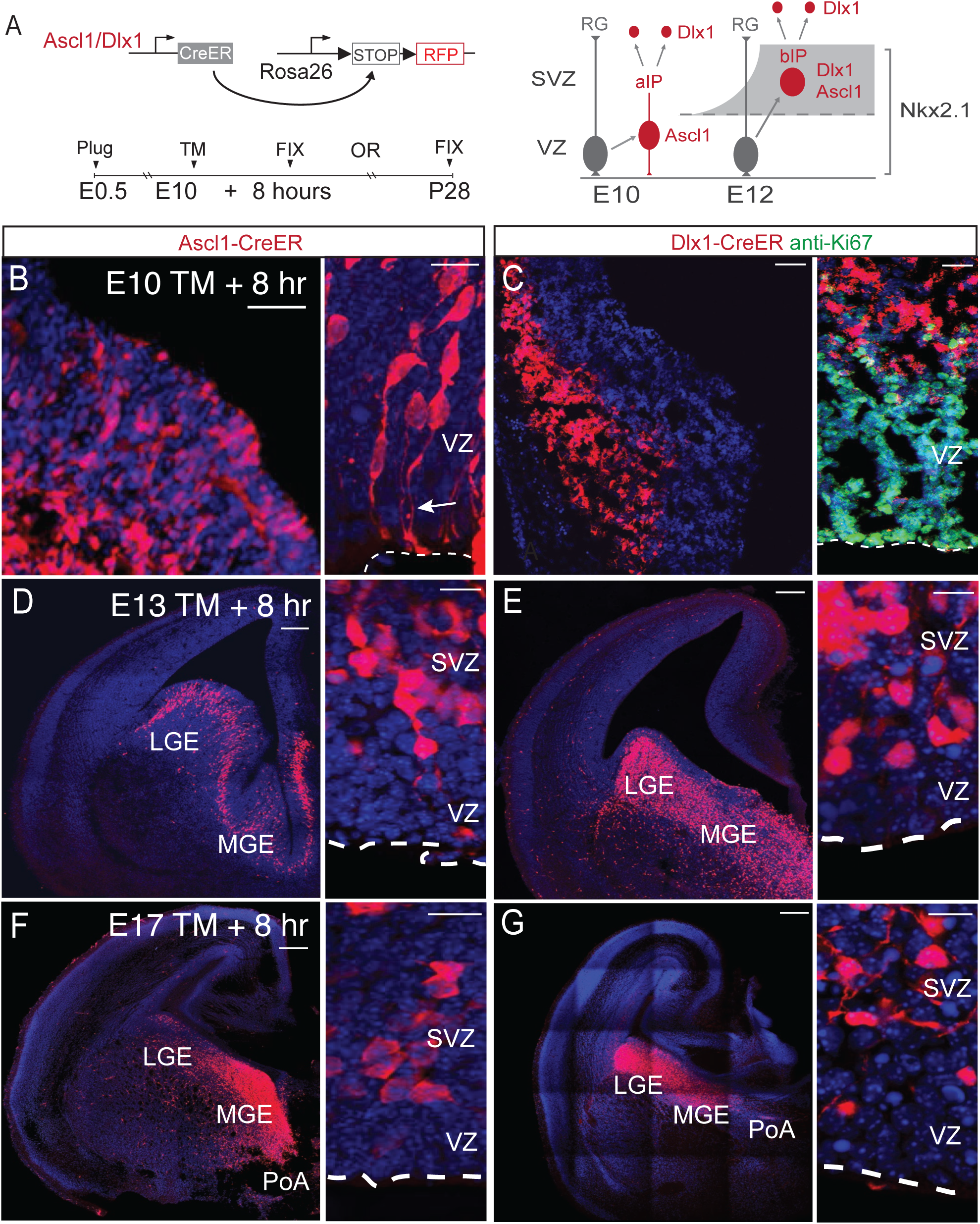
***Ascl1-CreER* and *Dlx1-CreER* pulse-chase distinguish aIPs and bIPs.** (A) Schematic of genetic fate mapping strategy using the *Ascl1* and *Dlx1* drivers. VZ: ventricular zone, SVZ: subventricular zone. (B) 8-hour pulse-chase in *Ascl1-CreER* following E10 TM induction labeled aIPs in MGE. Note apical endfeet (arrow) and absence of basal processes. (C) 8-hour pulse-chase in *Dlx1-CreER* following E10 TM induction labeled postmitotic neurons in MGE, which were negative for the mitotic marker Ki67. (D-G) 8-hour pulse-chase in *Ascl1-* or *Dlx1-CreER;Ai14* mice at E13 revealed that bIP*^Ascl1^* and bIP*^Dlx1^* were restricted to the SVZ (right subpanels) of the subpallium at E13 (B, C) and E17 (D, E). While the distributions of IP*^Ascl1^* and IP*^Dlx1^*overlapped closely at E13 (D, E), they diverge and have opposing dorsal-ventral gradients by E17 (F, G), with IP*^Dlx1^* concentrated more dorsally and IP*^Ascl1^* more ventrally within the MGE. Scale bars: B, C: 20 um left panels, 10 um right panels; D-G: 50 um left panels, 10 um right panels.

To specifically target IPs in the MGE and exclude populations from LGE and CGE, we used an intersectional strategy combining *Ascl1*-or *Dlx1-CreER* with the *Nkx2.1-Flp* driver and *Ai65* reporter (Figure 4A). Short 8-hour pulse-chase in *Ascl1-CreER;Nkx2.1-Flp;Ai65* mice demonstrated specific labeling of bIPs in the SVZ within the MGE but not LGE and CGE (Figure 4H-K). We then fate mapped interneuron progeny in P28 cortex derived from MGE IPs after TM induction across embryonic stages. E10 TM induction in *Ascl1-CreER;Nkx2.1-Flp;Ai65* mice yielded a sparsely-labeled and rather limited set of interneurons in L6, L5, L3 (Figure 4B-G). Compared with those fate-mapped from *Nkx2.1-CreER;Ai14* mice at the same stage (Figure 1P), the distribution of these interneurons had a more tightly L6 restricted laminar pattern, possibly due to these aIPs becoming exhausted soon after their divisions (Figure 4G). These results suggest that early aIPs are limited in number, cell division and fate potential, consistent with the fact that they derived from early RG^E^s, which have limited neurogenic capacity.

**Figure 4.**
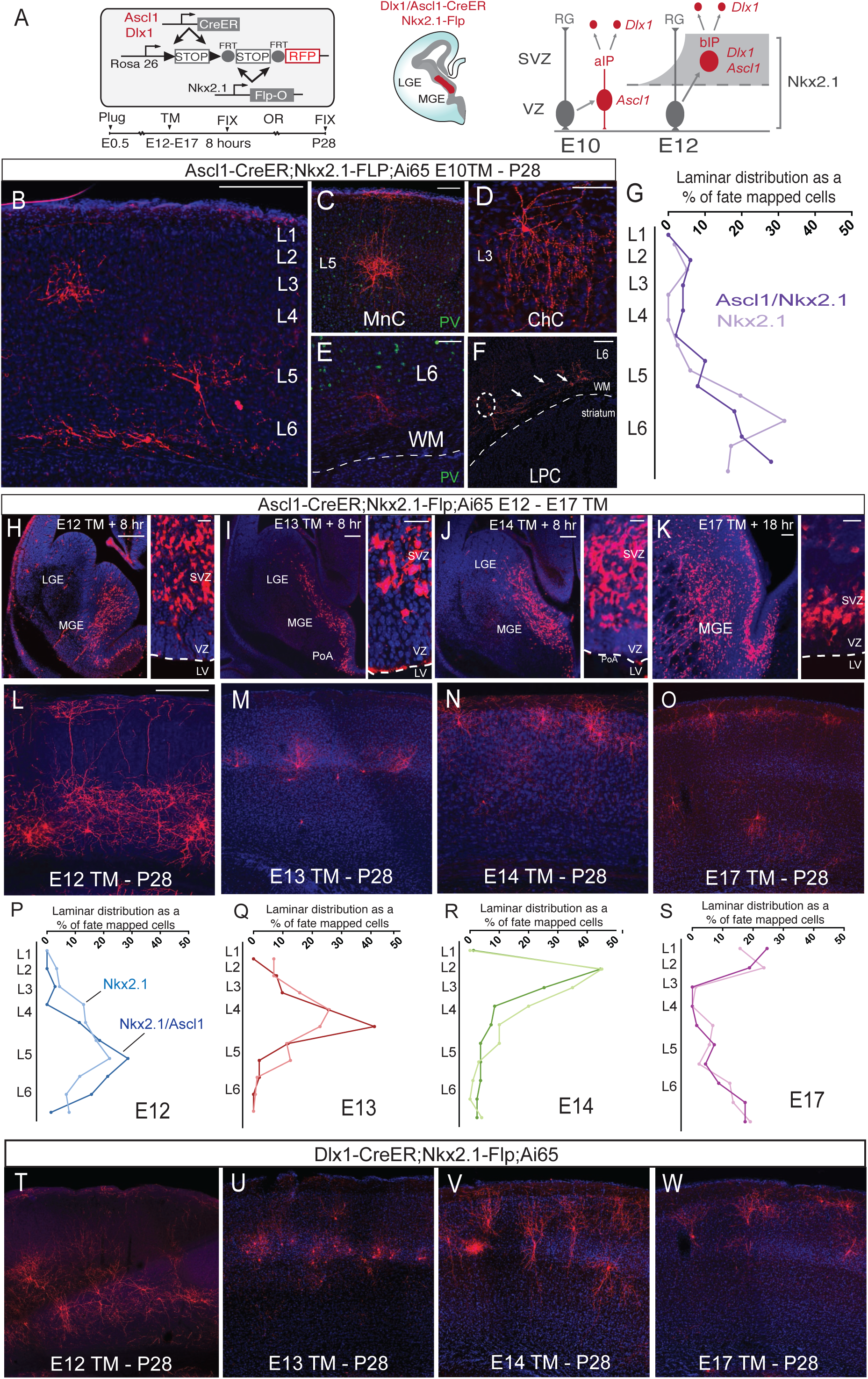
Fate-restricted Ascl1 and Dlx1 bIPs diversify and amplify interneuron production in an inside-outside-inside sequence. (A). Scheme of inducible and intersectional fate mapping of apical and basal intermediate progenitors expressing either *Ascl1* or *Dlx1* in the MGE (*Nkx2.1*^+^) between E10 and E17. (B-F) Intersectional *Nkx2.1*/*Ascl1* fate mapping following E10 TM induction allows resolution of several major GABAergic cell types in L3, L5, and L6 of P28 cortex. These include most notably L5 Martinotti cells with prominent axon projection to L1 (C; MnC), L2/3 ChCs with dense and local axon arbors characterized by vertically oriented cartridges of presynaptic boutons (D), and L6 GABAergic long-projecting cells (LPCs, E) with axons (dashed circle, arrows in F) extending in the white matter (WM). (G) The laminar distribution of GABAergic neurons fate-mapped following E10 TM induction in *Ascl1/Nkx2.1* (light purple) and *Nkx2.1* progenitors (dark purple) have overall similar profiles, with a sharper output of L6 interneurons from the likely more differentiated *Ascl1/Nkx2.1* progenitors. (H-K) Short pulse intersectional labeling of IPs *^Ascl1^* in *Ascl1-CreER;Nkx2.1-Flp;Ai65* mice on E12, E13, E14, and E17. Note the absence of labeling in adjacent LGE, in contrast to patterns in (Figure 3 D-G). (L-O) Fate mapping of IPs *^Ascl1^*at E12, E13, E14, and E17 reveal interneuron progeny with highly restricted laminar distribution and morphology in mature cortex. Note the strict inside-out pattern between E12-14 and reappearance of deep layer cells (arrow) in fate mapping at E17.5 (O), completing the inside-out-inside sequence. (P-S) Each temporal cohort of IPs*^Ascl1^* produces a set of more laminar-restricted interneurons (darker lines) than those fate-mapped from *Nkx2.1^+^* progenitors, which include both RG and IPs (lighter lines) from the same embryonic date. Note the re-appearance of deep layer cells in E17.5 fate mapped cortex (S). The temporal outputs of IPs*^Ascl1^* demonstrate a clear inside-out-inside sequence along the progression of the RG*^L^* lineage. (T-W) Fate mapping of IPs*^Dlx1^* at E10, E12, E13, E14, and E17 in *Dlx1-CreER;Nkx2.1-Flp;Ai65* mice labels interneuron subsets with similar laminar distribution but different morphological features compared to those fate-mapped from IPs*^Ascl1^*. See Figure S3 for further morphological characterization. Scale bars: B: 300 um; C, D, E, F: 50 um; H-K: 50 um in left panels, 10 um in right panels; L-O: 300 um; T-W: equivalent scale to L-O.

From E12 to E17, there was a dramatic change in the profile of interneuron production from bIPs compared to aIPs, both in terms of cell numbers and their laminar patterns (Figure 4H-W). First, equivalent TM doses consistently labeled a much larger number of interneurons than was observed for E10 induction. Second, at consecutive embryonic days, TM induction labeled a much more laminar-restricted set of interneurons compared to those from same day induction in *Nkx2.1-CreER;Ai14* mice (Figure 4 L-S), suggesting that bIPs are much more restricted in their laminar fate potential. Third, between E12-E17, bIP*^Ascl^*-derived interneurons laminated the cortex in an inside-out-inside sequence (Figure 4 L-S). Fourth, equivalent fate mapping experiments in *Dlx1-CreER;Nkx2.1-Flp;Ai65* mice between E12-17 revealed that bIPs*^Dlx1^* generated interneurons that laminated in an inside-out sequence, without the last set of deep layer interneurons (Figure 4T-W). Together, these results suggest that, from the lineage progression of RG^L^s, large numbers of fate-restricted bIPs are generated as temporal cohorts that amplify interneurons destined to specific cortical layers in an inside-out-inside sequence.

### Multiple temporal cohorts of bIPs diversify interneuron types

In addition to laminar position, cortical interneurons consist of dozens of cell types distinguished by multi-modal features including morphology and marker expression (Huang and Paul 2019; Tremblay et al. 2016). To explore the relationship between IPs and interneuron types, we characterized multiple morphological and marker-defined interneuron types across IP-mediated neurogenesis, using low dose TM induction to sparsely label interneuron in adult cortex.

Although early aIPs (fate mapped at E10 in *Ascl1-CreER;Nkx2.1-Flp;Ai65* mice) gave rise to a rather limited number of interneurons (Figure 4B-G), the progeny nonetheless included multiple distinct morphological types. For example, the L6 cells that extended axons into the white matter were morphologically long projecting GABAergic neurons (Figure 4F). Additionally, L5 Martinotti cells (MNCs) could be identified by their prominent L1 axons, (Figure 4C), distinguishable from PV-positive L5 large basket cells (Figure 4E). Surprisingly, early aIPs also consistently generated a small number of clearly identifiable L2/3 ChCs (Figure 4D), indicating that ChCs are not all generated during the late phase of MGE neurogenesis as previous studies suggested (Sultan et al. 2018; Taniguchi et al. 2013). Therefore, we find that early aIPs generate a limited yet distinct set of cortical GABAergic neuron types.

From E12 onward, a much larger set of morphologically diverse interneuron types were systematically and orderly generated from bIPs across embryonic times. At E12, interneurons fate-mapped from bIPs*^Ascl1^*were largely restricted to the production of L5 cells including MNCs, PV^+^ large basket cells (LBSKs), and large dense arbor cells (LDACs) (Figure 4H,L, Figure S3A-B). E13 fate-mapped cells mainly resided in L4 and included a set of highly characteristic locally projecting ‘bushy’ cells, small basket cells (SBSKs) and non-Martinotti cells (non-MNCs) (Figure 4I,M, Figure S3A, B). E14 fate-mapped cells mainly localized to L2/3 and included large sets of L3 MNCs, non-MNCs, BSKs and LDACs (Figure 4J,N, Figure S3A, B). Importantly, we observed that E17 fate-mapped cells became essentially bimodal in terms of their laminar location, with one cohort residing in L2 near the L1 border and another in L6 and deep L5. Moreover, we found that E17 progenitors produced a rich population of ChCs and were otherwise binary in producing only two other distinct morphological types: horizontal arbor cells (HACs) in L2 and L6, and descending arbor cells (DSCs) in L2 (Figure 4K,O; Figure S3A-E). ChC, HAC and DSCs together accounted for greater than 90% of the total E17 fate-mapped cells in three cortical areas examined (somatosensory, motor, and medial prefrontal cortex; Figure S4B-D). Therefore, at successive stages of MGE neurogenesis, multiple temporal cohorts of bIPs*^Ascl1^* are each become dedicated to generate sets of interneurons with distinct features and laminar identity in an inside-out-inside sequence.

Similar to experiments with *Ascl1-CreER*, fate mapping with *Dlx1-CreER;Nkx2.1-Flp;Ai65* mice showed that bIPs*^Dlx1^*also generated diverse and sharply layer-restricted sets of interneuron types. Between E12-E14, bIPs*^Dlx1^*-derived interneuron types were overall similar to those from bIPs*^Ascl1^*, consistent with the possibility that most bIPs during this period expressed both *Ascl1* and *Dlx1* (Figure 4T-V, Figure S3A,B). Thereafter, interneuron progeny from bIPs*^Dlx1^* and bIPs*^Ascl1^* became more distinct in ratio and types. In particular, there were rather few L2 ChCs, L2 HAC, DSC or L5/6 ChCs generated from E16-17 bIPs*^Dlx1^* (Figure 4W; Figure S3A, B). Of note, it is possible that the small numbers of bIPs*^Dlx1^* that did give rise to ChCs at this time may also express *Ascl1*. Conversely, there were few BSKs and MNCs generated from E16-17 bIPs*^Ascl1^*in the examined areas of cortex (Figure S3A). Together, these results suggest that, along the course of MGE neurogenesis, multiple temporal cohorts of fate restricted IPs diversify as well as amplify cortical interneuron types, with early aIPs and later bIPs likely reflecting the temporal progression of RG^E^s and RG^L^s, respectively.

### Two separate waves of ChCs are generated toward the end of RG^E^ and RG^L^ neurogenesis

As ChCs appear to be the last set of interneurons generated from both the RG^E^ and RG^L^ lineages and can be most reliably identified, we systematically characterized the generation of this cell type to further delineate the temporal relationship between RG^E^-and RG^L^-mediated of neurogenesis. We carried out a comprehensive morphological screen in both *Nkx2.1-CreER;Ai14* and *Ascl1-CreER;Nkx2.1-Flp;Ai65* fate mapping cohorts to characterize the generation of supragranular and infragranular ChCs across embryonic neurogenesis.

E10 TM induction in *Nkx2.1-CreER;Ai14* mice consistently labeled a set of supragranular layer ChCs (Figure 1P, 5A, B). Although overall only a small number of upper layer interneurons were labeled, up to 75% of these were ChCs (Figure 5E). Interestingly, a significant portion of these E10 fate-mapped ChCs resided in L3, extended more vertical dendrites toward the pia, and elaborated axon arbors within L3 and 4 (Figure 5B); these characteristics are in contrast to the typical L2 ChCs, which resided at L1/2 border, extended wide-angled layer 1 dendrites, and elaborated axon arbors restricted to L2/3 (Figure 6, Figure S4A). We then quantified the soma depth of all upper layer ChCs labeled from E10 to E17 TM induction in both *Nkx2.1-CreER* and *Ascl1-CreER;Nkx2.1-Flp* mice. This analysis revealed that whereas L2/3 ChCs were labeled by E10 and E12 induction in *Nkx2.1-CreER;Ai14* mice (inductions that target both RGs and IPs), ChCs were instead labeled from E12 and E13 induction in the *Ascl1-CreER;Nkx2.1-Flp;Ai65* mice (such inductions target mostly bIPs) (Figure 5C). These results suggest that the early set of L2/3 ChCs were likely generated from RG^E^s predominantly through *Ascl1^+^* aIPs around E12 and E13. Indeed, BrdU birth dating confirmed that these ChCs, designated as ChCs^E^, were born at approximately E13 (Figure 5D).

**Figure 5.**
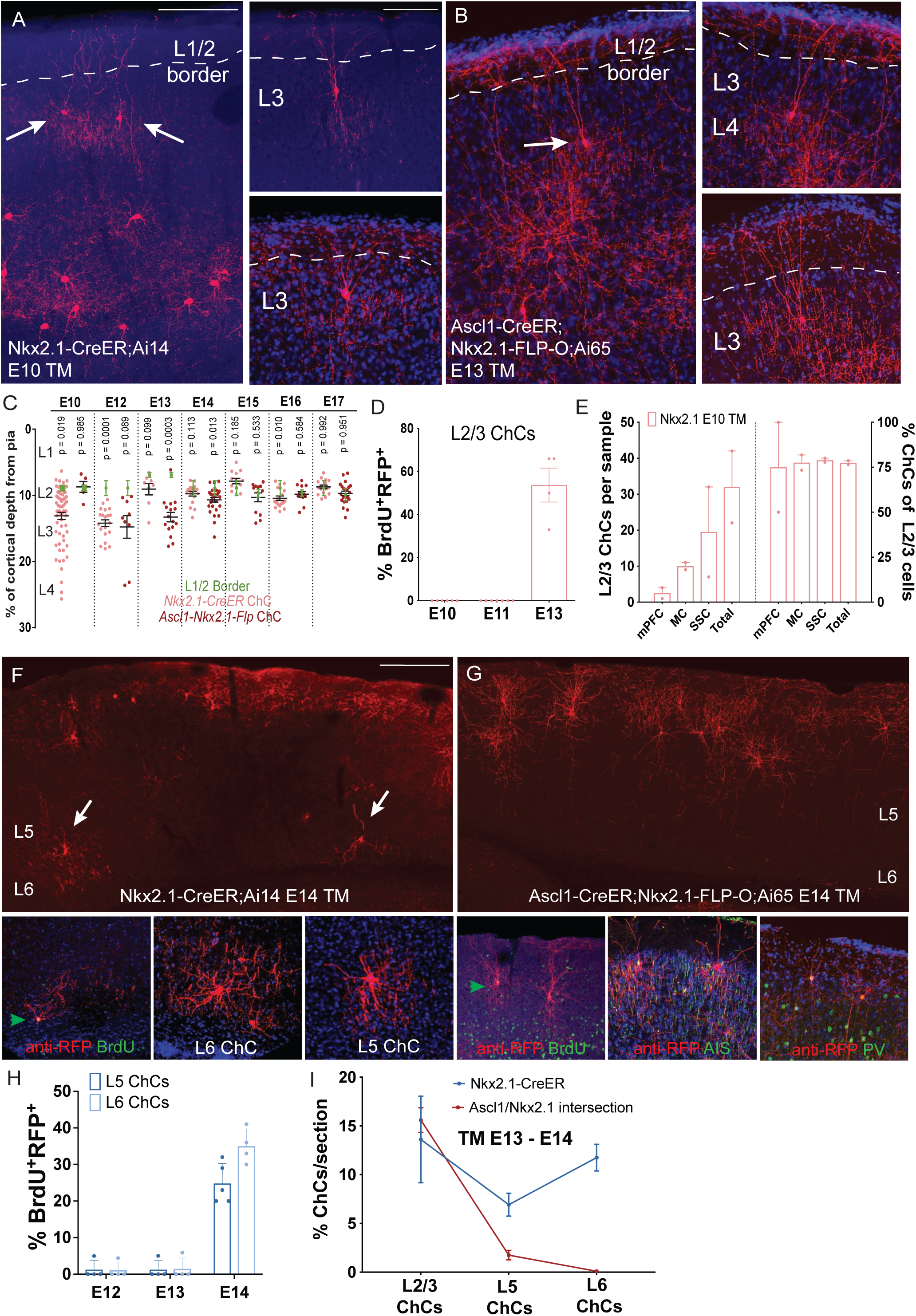
A small first wave of ChCs is generated from RG^E^s and laminates in an outside-in order. (A) E10 TM induction in *Nkx2.1-CreER;Ai14* mice labels an early set of L3 ChCs (arrows), which localized well below the L1/2 border (dashed lines), extended dendritic branches toward the pia, and elaborated axon arbors in L3 and L4. Scale bar: 300 um in left panel, 100 um in right panel. (B) E13 induction in *Ascl1-CreER;Nkx2.1-Flp;Ai65* mice labeled L3 ChCs that resembled those in (A), suggesting that they were born on E13 likely from *Nkx2.1^+^* RG^E^s present at E10 that self-renewed for several days. Scale bar: 100 um in left panel, right panel equivalent scale to (A, right). (C) Quantification of the cortical depth distribution of upper layer ChCs in MC and SSC labeled through fate mapping of both *Nkx2.1^+^*(pink dots) and *Ascl1^+^/Nkx2.1^+^* progenitors (maroon dots) from E10 to E17. Early RG^E^-derived L2/3 ChCs are followed by a large second wave of later-born L2 ChCs strictly localized to the layer 1/2 border, p values were derived from a T-test comparing the mean depth of cells at the layer 1/2 border (green) to the mean depth of ChCs from that genotype and time point (black). (D) The early set of L2/3 ChCs derives from the late phase of RG*^E^* cell division, as shown by an increased rate of BrdU incorporation on E13, but not on E10 or E11. (E) A large proportion (∼75% across mPFC, MC and SSC) of L2/3 cell progeny generated from RG^E^s are ChCs (cell counts plotted on left y-axis, proportion of cells plotted on right y-axis) (F, G) First-wave L5/L6 ChCs are produced from E14 *Nkx2.1^+^*RGs (F), but not from *Ascl1^+^/Nkx2.1^+^* IPs (G), which instead produce only L2/3 ChCs and other L2/3 interneurons. ChCs were confirmed by their innervation of pyramidal neuron AIS (marked by ankyrin G), birthdated with BrdU, and distinguished from PV basket cells with parvalbumin (bottom subpanel in G). Scale bar: F, G: 300 um in top panels, scale in lower subpanels comparable to (A, right). (H) The early set of L5/6 ChCs fate-mapped from E12 TM induction incorporated BrdU almost exclusively on E14, rather than E12 or E13, suggesting these cells are born on average one day later than L2/3 ChCs from RG^E^s. (I) Quantification of the percentage of L2/3, L5 and L6 ChCs fate-mapped from *Nkx2.1^+^* (blue) or *Ascl1^+^/Nkx2.1^+^* (red) progenitors from E13 to E15. Note that the early set of L5/6 ChCs were not generated from *Ascl1^+^/Nkx2.1^+^* IPs and thus were likely generated directly from *Nkx2.1^+^* RG^E^s.

**Figure 6.**
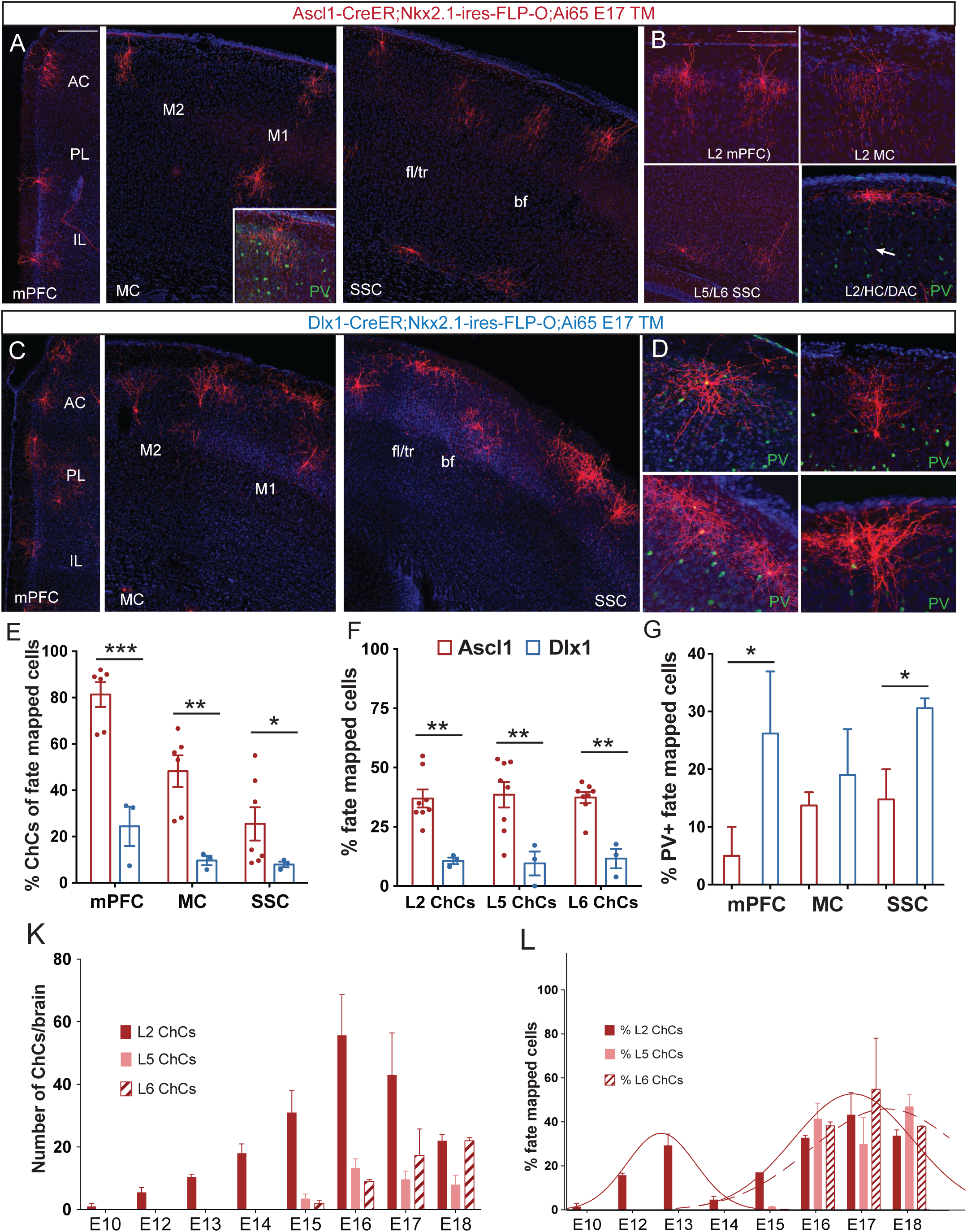
A large second wave of ChCs is generated from bIP and laminate in an outside-in order. (A, B) At E17, IPs*^Ascl1^* produce a large set of L2, L5, and L6 ChCs and HC/DACs. Late-born ChCs are often weakly PV-immunopositive (green). AC: anterior cingulate; PL: prelimbic, PL; IL: infralimbic; M2: secondary motor; M1: primary motor; fl/tr: forelimb/trunk somatosensory; bf: barrel field. Scale bar, 100 um in A, B. (C,D) At E17, IPs*^Dlx1^* generate primarily L2/3 non-ChC interneurons including a large number of PV^+^ basket cells in L2/3 (D). Scales equivalent to A, B (E) IPs*^Ascl1^* (red) generate a significantly more pure population of ChCs in mPFC, MC and SSC relative to IPs*^Dlx1^* (blue) from E16-E18, quantified as the relative proportion of total fate-mapped cells in each genotype. (F) All three laminar ChC subtypes (L2, L5, L6) are produced at a much higher rate from IPs*^Ascl1^* (red) relative to IPs*^Dlx1^*(blue) from E16-E18. (G) IPs*^Dlx1^* (blue) generate significantly more PV basket cells (PV^+^ ChCs excluded) relative to IPs*^Ascl1^* (red) during the same period, quantified as the relative proportion of total fate-mapped cells in each genotype. (K) Daily output of ChC laminar subtypes from *Ascl1^+^/Nkx2.1^+^* progenitors throughout MGE neurogenesis, quantified as number of ChCs summed from mPFC, MC and SSC. (L) Quantification of the relative proportion of ChC laminar subtypes fate-mapped from *Ascl1^+^/Nkx2.1^+^* progenitors reveals two distinct waves for ChC production via IPs. The first wave consists of mainly of L3 ChCs generated from RG*^E^* and aIPs. The second wave consists of L2, L5 and L6 ChCs produced from RG*^L^* via bIPs*^Ascl1^*. The two waves of ChC production closely fit two sequential normal Gaussian distributions. Solid curves reflect nonlinear fit to L2/3 ChC data only with R^2^ value of 0.7866 (left) and 0.8431 (right) and D’Agostino & Pearson omnibus normality test P value 0.0029 (left) and 0.0168 (right). Dashed curve reflects non-linear fit to L2, L5 and L6 ChCs combined with R^2^ value of 0.6954 and D’Agostino & Pearson omnibus normality test P value 0.0952.

Within E14-induced *Nkx2.1-CreER;Ai14* mice there was a small set of L5/6 ChCs (Figure 5F). Combined TM and BrdU labeling at E14 confirmed that these deep layer ChCs were born at this time (Figure 5H). Furthermore, in experiments with E12 TM followed by BrdU pulses at E12, E13, or E14, we observed that deep layer ChCs were only co-labeled by the E14 BrdU pulse (Figure 5I). This result suggested that these L5/6 ChCs were generated after L3 ChCs (Figure 5D) from RG^E^s. Notably, although RG^L^ and bIPs also mediate active neurogenesis at E14, they almost exclusively generated supragranular interneurons at those times (Figure 1S, Figure 4N,R). Indeed, TM induction at E14 in *Ascl1-CreER;Nkx2.1-Flp;Ai65* mice did not label any deep layer ChCs (Figure 5G, I). Together, these results suggest that the E14-born deep layer ChCs might be generated directly from *Nkx2.1^+^* RG^E^s at the end of their lineage.

Earlier-born L2/3 ChCs were followed by a much larger set of late-born L2 ChCs, which were almost always found to be strictly located at the L1/2 border (Figure 6A, B). These late ChCs, designated as ChCs^L^, derived from fate mapping cohorts of both *Nkx2.1-CreER* and *Ascl1-CreER;Nkx2.1-Flp* mice. While most are likely generated from bIPs*^Ascl1^* given the dominance of this progenitor type in MGE neurogenesis at this time, we cannot exclude the possibility that some may derive directly from *Nkx2.1^+^*RGs. However, TM induction at E16-E18 in *Ascl1-CreER;Nkx2.1-Flp;Ai65* mice in particular generated a much larger and purer set of infragranular layer ChC^L^s than any other fate mapping paradigm, also supporting the idea that bIPs ^Ascl1^ were the dominant population responsible for ChC production in the final stage of embryonic development (Figure 6A, B, E-G). In comparison, bIPs^Dlx1^ generated rather few ChCs^L^ (Figure 6C-G), and it is even possible that ChCs^L^-generating bIPs^Dlx1^ may in fact be Ascl1^+^, while Ascl1^-^/Dlx1^+^ bIPs might not generate ChCs^L^ at all.

Quantification of the proportion of ChCs normalized to total fate-mapped cells produced in *Ascl1/Nkx2.1* intersection over time revealed two sequential waves of L2/3 ChC production, peaking at ∼E13 and ∼E17, respectively. Each wave fit a normal Gaussian distribution following a nonlinear regression analysis (Figure 6K, L). Therefore, supragranular layer ChCs are likely generated in two waves. First a small set is produced by RG^E^ and/or aIPs between E12-E13, followed by a second overlapping and larger set, mostly via bIPs*^Ascl1^* between E15-E17. Together, these results provide strong evidence that two separate sets of ChCs are generated toward the end of the two consecutive waves of MGE neurogenesis outlined herein, each with an outside-to-inside lamination pattern. While the RG^E^ wave generates a small set of L2/3 and L5/6 ChC^E^s and completes by ∼E14, the RG^L^ wave generates a much larger set of L2 and L5/6 ChC^L^s and completes at ∼E17.

## DISCUSSION

### The progenitor and lineage basis of MGE interneuron generation and diversity

Recent large-scale single-cell RNA sequencing has unveiled a working draft of cortical GABAergic neuron taxonomy across the mammalian species (Tasic et al. 2018; van Velthoven et al. 2025; Schmitz et al. 2022; Bakken et al. 2021) and have suggested a molecular basis of interneuron type definition based on synaptic communication styles (Paul et al. 2017; Huang and Paul 2019). However, the developmental basis of this spectacular interneuron diversity remains not well understood. Despite major progress in understanding subpallial progenitor domains marked by transcription factor expression (Hu et al. 2017; Kepecs and Fishell 2014; Sultan and Shi 2018; Shi et al. 2021; Bandler and Mayer 2023), multiple progenitor types (Turrero García and Harwell 2017), and the temporal features of neurogenesis on interneuron production (Hu et al. 2017; Bandler et al. 2017; Bright et al. 2025), we have yet to achieve a cellular resolution lineage framework linking progenitors and their lineage progression to diverse interneuron types.

Combining intersectional fate mapping of progenitor types and their interneuron progeny across the full course of MGE neurogenesis, our results delineate a lineage framework in the MGE whereby multipotent RGs deploy distinct temporal cohorts of fate-restricted IPs to sequentially amplify and diversify interneuron types. According to this model (Figure 7), two separate pools of multipotent RGs mediate consecutive waves of neurogenesis, each deploying distinct sets and temporal cohorts of IPs to sequentially generate different sets of interneuron types that laminate the neocortex in an inside-out-inside order. Following the formation of MGE at ∼E9.5 marked by the onset of NKX2.1 expression (Sussel et al. 1999), NEs and RGs may consist of subpopulations with low or higher levels of Nkx2.1 expression (Figure S1A,D,G)). Whereas Nkx2.1^hi^ progenitors initiate early neurogenesis as RG^E^s captured by fate mapping in *Nkx2.1-CreER;Ai14* mice at E10, Nkx2.1^lo^ progenitors may largely evade *Nkx2.1-CreER* fate mapping at this time point and continue to amplify or remain quiescent, instead initiating neurogenesis at a later stage. RG^E^s are restricted to the caudal MGE, have limited neurogenic capacity, and involve *Ascl1^+^Dlx1^-^* aIPs that generate a small set of interneurons. These include L6 LPCs, L5 MNCs and PV LBCs, L2/3 ChCs, and L5/6 ChCs, generated between E10-E14 and laminating the cortex in an inside-out-inside sequence. Then, from about E12 onward, a large set of Nkx2.1 RG^L^s initiate neurogenesis throughout the full rostral-caudal axis of MGE with expanded neurogenic capacity. These RG^L^s leverage sequential temporal cohorts of *Ascl1^+^* and *Dlx1^+^*bIPs to generate a massive number and diverse set of interneurons which also laminate the cortex in an inside-out-inside sequence, including most if not all major morphological types of MGE interneurons. As RG^E^s and RG^L^s differ in spatiotemporal progression patterns, neurogenic capacity, IP types, and interneuron outputs, we suggest that they represent separate RG pools and mediate two consecutive (though partially overlapping) waves of neurogenesis rather than two stages of the same process of MGE neurogenesis. This scheme may reconcile two seemingly inconsistent conclusions on MGE lineages drawn from two previous retroviral fate mapping studies of (Ciceri et al. 2013; Sultan et al. 2018). While fate mapping at E11.5 (Ciceri et al. 2013) would have captured the RG^E^s with similarly limited capacity and laminar output patterns, fate mapping at E12.5 (Sultan et al. 2018) mostly likely have captured the RG^L^s with large capacity and broad laminar output patterns.

**Figure 7.**
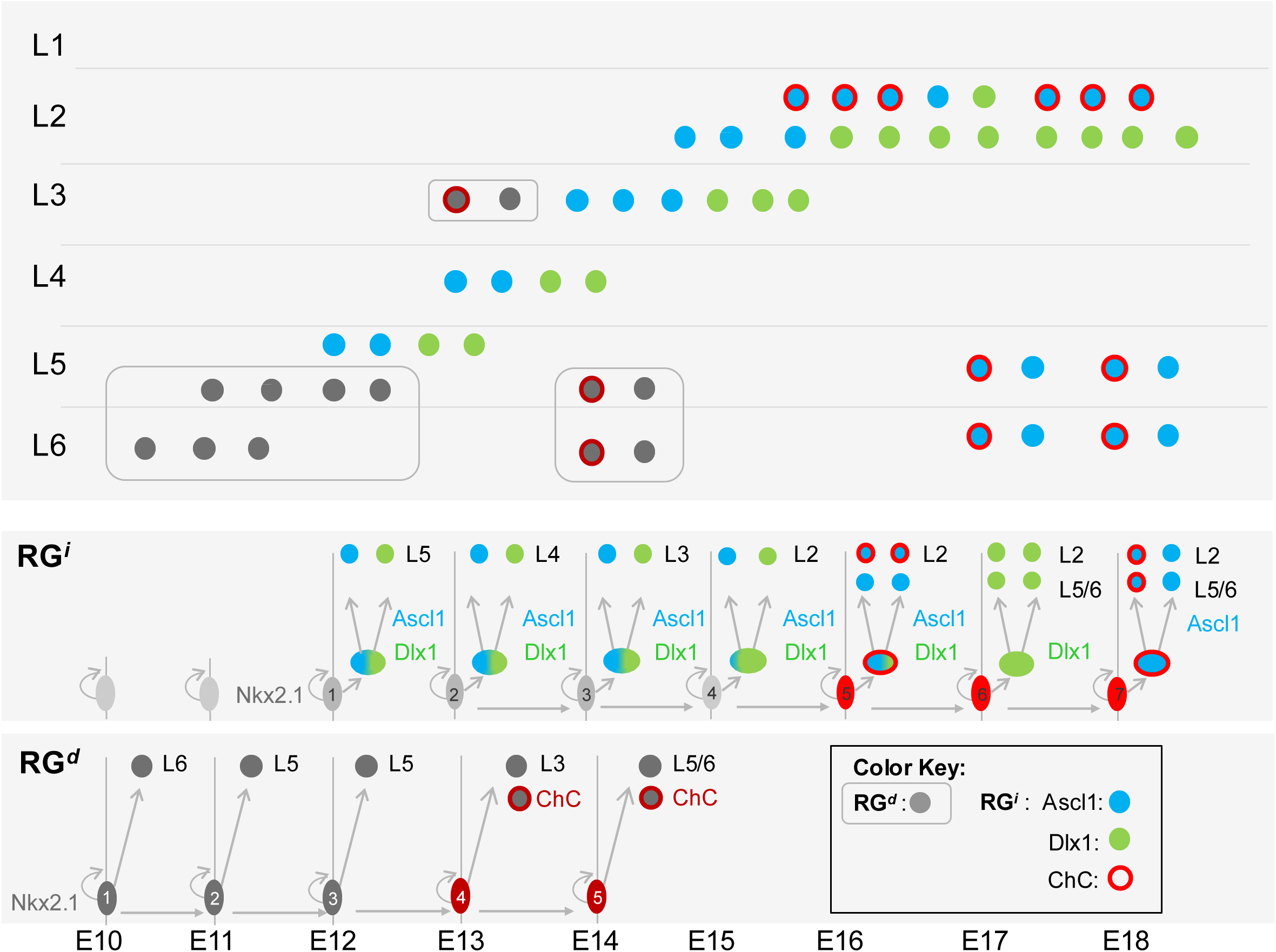
Multipotent RGs and fate-restricted IPs sequentially generate diverse cortical interneuron types. (A) Schematic laminar distribution of the early (E) and later (L) waves of MGE-derived interneurons. Colors are matched with morphological types listed at the bottom. LPC, long projection cell; MNC, Martinotti cell; BSK, basket cell; n-MNC, non-Martinotti cell; ST, stellate cell; HC, horizontal cell; ChC, chandelier cell. (B) A model on the progenitor and lineage mechanism of the generation of diverse interneuron types. aIP and bIP^A^ are distinguished by their *Ascl1* and *Dlx1* expression. bIP^A^ may later diverge to bIP^B^ and bIP^C^, which express either *Ascl1* or *Dlx1* and are likely present in parallel in the later phase of RG^L^s progression. See Discussion text for detailed description.

In addition to the rostral-caudal axis, the dorsal-ventral axis of MGE is marked by TF expression subdomains (Flames et al. 2007; Long et al. 2009). It is possible that the dorsal-ventral axis may comprise additional multi-potent RG pools with different fate potentials, which generate different compositions and/or ratios of interneuron types. Testing this hypothesis may require new methods for large scale clonal analysis with cellular resolution.

### Temporal cohorts of fate-restricted bIPs amplify and diversify interneuron production

Although IPs are thought to mediate the bulk of MGE neurogenesis and several types of IPs have been found (Brown et al. 2011; Pilz et al. 2013; Turrero García and Harwell 2017), their precise roles in the amplification and diversification of interneurons had not been examined directly due to the lack of specific fate mapping tools. A previous study suggested that indirect neurogenesis from IPs is biased for the production of PV cells while direct neurogenesis from RGs is biased for SST cells (Petros et al. 2015). Here, we have achieved specific and systematic fate mapping of MGE IPs as defined by Ascl1 expression using an intersectional strategy. We found that, between E10-E12, *Ascl1^+^*IPs were a rather small population of aIPs associated with RG^E^s; they were *Dlx1^-^* and generated a small set of interneuron progeny. In sharp contrast, from E12 onward, the number of *Ascl1^+^*IPs massively expanded, which were exclusively bIPs associated with RG^L^s. bIPs likely consisted of *Ascl1^+^*/*Dlx1^+^*, *Ascl1^+^Dlx1^-^, Ascl1^-^ Dlx1^+^* subpopulations. These temporal cohorts of bIPs were fate-restricted to generate laminar specific subset of interneurons. Together, these results suggest that the generation of temporal cohorts of fate-restricted bIPs is a mechanism whereby multi-potent RGs amplify and diversify interneuron production along their lineage progression. The extent of the fate restriction of individual IPs, e.g. whether generating a pair of cells of the same morphological type, (e.g. ChCs), remains unknown.

### The outside-in pattern of ChCs lamination suggests cell intrinsic laminar identity

Among diverse cortical interneurons, the axo-axonic ChCs can be unequivocally recognized as a bona-fide type based solely on their distinct morphological features (Raudales et al. 2024; Somogyi et al. 1982; Woodruff et al. 2010). Our previous fate mapping suggested that ChCs are generated from the late embryonic stage (Taniguchi et al., 2013). More recent retroviral-mediated studies convincingly demonstrated that laminar subsets of ChCs are generated toward the end of RG lineage progression in an outside-sequence (Sultan et al. 2018). Here, by fate mapping across the full time course of MGE neurogenesis, we found that in addition to a large set of late-born ChCs, a small yet significant set of ChCs are generated at a surprisingly early time between E12-13. While the late-born ChCs are generated toward the end of the RG^L^ lineage through bIPs, the early set is generated toward the end of RG^E^. Both sets of ChCs settle in the cortex in an outside-in sequence.

Laminar subsets of ChCs have been described in multiple cortical areas and species (Inda et al. 2009; Lewis and Lund 1990; Somogyi et al. 1982; Raudales et al. 2024). Complete single cell reconstruction and analysis of dendrite-axon distribution suggest that these laminar subsets may have distinct input-output connectivity patterns (Wang et al. 2019). However, the developmental basis of the laminar subtypes of ChCs is unclear. It is possible that post-mitotic ChCs are unspecified in terms of their laminar destination, and subsequently acquire this identity during migration through extrinsic signals (e.g. cell interactions, morphogens). However, a previous transplantation experiment suggests that laminar settlement of late-born ChCs is likely determined by cell intrinsic properties (Taniguchi et al. 2013). Here we substantiate this conclusion by showing that both early-and late-born cohorts of ChCs laminate in an outside-in sequence. In particular, E12-born ChCs specifically settle in L2/3 and a large fraction of late-born ChC target deep layers - a laminar deployment pattern opposite to that of other MGE interneurons born at the same time. These results suggest that young ChCs do not follow the generic cues for MGE interneuron migration but are destined to specific laminar positions; thus laminar deployment of ChCs could be largely guided by cell intrinsic information. The finding that L2/3 ChCs are generated at early (∼E12) as well as late (∼E15) stages of MGE neurogenesis further raises the questions of whether they differ in the schedule of their migration and laminar settlement, and in their connectivity to subsets of pyramidal neurons

### The developmental basis and evolutionary implication of two spatiotemporal RG pools

What is the developmental logic of two separate MGE RG pools with different spatiotemporal characteristics and neurogenic mechanisms? As RG^L^s have a much larger capacity, why are RG^E^s necessary? We suggest that the distinction and spatiotemporal separation of RG^E^s and RG^L^s might be rooted in the evolutionary history of MGE/POA subdivisions in light of the expression and function of the homeodomain transcription factor (TF) OXT2 (Acampora et al. 2005).

During the developmental patterning of vertebrate telencephalon, OTX2 is a foundational TF that maintains forebrain identity after its specification. OTX2 is expressed in the rostral patterning center (RPC), ventral telencephalon (subpallium, ganglionic eminence – GE), and caudodorsal telencephalon (Acampora et al. 2005; Simeone et al. 2002). Within the GE, OTX2 is expressed throughout the MGE and POA largely in parallel with NKX2.1 (Hoch et al. 2015). Importantly, OTX2 plays a key role in regulating the regional subdivision of MGE and in promoting MGE neurogenesis (Hoch et al. 2015). OTX2 is necessary to maintain the rostral and ventral characteristics of MGE by repressing POA and caudal MGE fates. Within the rostral-ventral MGE, OTX2 promotes neurogenesis specifically in the SVZ (i.e. intermediate progenitors) by repressing the anti-neurogenic factors (Hes1, Id4), thereby enhancing neurogenic factors ASCL1 and DLX1 (Hoch et al. 2015). These findings highlight the correspondence between OTX2 function in the MGE with the spatiotemporal division of RG^E^s and RG^L^s. With its higher expression in the SVZ and its role in repressing anti-neurogenic genes in SVZ (Hoch et al. 2015), it is possible that OTX2 may primarily function in IPs, which are much expanded in the RG^L^s lineage as bIPs. As OTX2 is required for rostroventral but not caudodorsal MGE identity (Hoch et al. 2015), RG^L^s and RG^E^s might involve different transcription programs and OTX2 might selectively promote the expansion and neurogenesis of RG^L^s over RG^E^s.

The basic organization of the subpallium, including the LGE, MGE and POA, may have its evolutionary origins in early vertebrates (Medina et al. 2014; Sugahara et al. 2017, 2016). Single cell transcriptome analyses in amniotes reveal that the number and diversity of pallial GABAergic interneurons are more limited compared to mammals (Tosches et al. 2018; Hain et al. 2022). Throughout vertebrate evolution, the predominant mode of neuron production is direct neurogenesis from RGs; indirect neurogenesis through IPs is found to a limited extent only in several species of reptiles and birds (through see (Deryckere et al. 2025)) and has vastly expanded in mammals (Martínez-Cerdeño et al. 2016). It is possible that the generation of cortical GABAergic neurons in the more ancient MGE/POA in reptiles is mediated by RGs that resemble RG^E^s in the caudal MGE in mice, with limited capacity and diversity. It is further possible that the rostral expansion of the mammalian telencephalon, in part regulated by Otx2, included the rostral MGE. This evolutionarily more recent MGE region, characterized by its innovation of RG^L^s and the expansion of bIPs constituting the prominent SVZ, may give rise to the much expanded number and diversity of mammalian neocortical GABAergic neurons, and set the stage for the vast expansion and further innovation of the human MGE SVZ neural progenitors (Jia et al. 2026).

## MATERIALS AND METHODS

### EXPERIMENTAL MODEL AND SUBJECT DETAILS

#### Mouse models

The following mouse strains were used for fate-mapping experiments studying GABAergic cortical interneurons: Nkx2.1-CreER (Taniguchi et al., 2011), Ascl1-CreER (Kim et al., 2011), Dlx1-CreER (Taniguchi et al., 2011), Nkx2.1-ires-FLP-O (He et al., 2016), Ai14 (Madisen et al., 2010), Ai65 (Madisen et al., 2015). Breeder mice used in fate mapping experiments were backcrossed 5 generations to Swiss-Webster background for genetic background homogeneity and due to these backgrounds having reduced susceptibility to side effects from tamoxifen dosing in these strains. All mouse colonies were maintained in accordance with protocols approved by the IACUC at Cold Spring Harbor Laboratory. Mice were housed by gender in groups of 2-4 with access to food and water ad libitum and a 12-hour light-dark cycle. Datasets for all fate-mapping experiments were assembled from groups of a minimum of two litters for each experiment, after processing multiple male and female brains processed from each. For all experiments, no relevant morphological or cellular composition differences were observed between male and female brains.

### METHOD DETAILS

#### Inducible genetic fate mapping

Nkx2.1-CreER, Ascl1-CreER, or Dlx1-CreER mice were bred separately to *Ai14* reporter mice to label MGE-derived interneurons and progenitors in single driver fate-mapping experiments. Mouse brain tissue was harvested for histology either during embryonic development or on P28 according to the inducible fate-mapping protocols described below. Similarly, to perform intersectional fate mapping of interneurons, Ascl1-CreER or Dlx1-CreER mice were bred with double-transgenic Nkx2.1-ires-FLP-O;Ai65 mice. In such experiments, the Ai65 reporter labeled cells with tdTomato only when both the lox-Stop-lox and frt-STOP-frt cassettes were excised by Cre recombinase and flippase within a single cell expressing both genetic drivers.

Embryos were staged in days post coitum, with embryonic day (E) 0.5 defined as noon of the day a vaginal plug was detected after overnight mating. Adult mice in fate mapping experiments were anesthetized with 2.5% avertin before transcardial perfusion with 4% paraformaldehyde (PFA) on postnatal day (P) 28. Brains were post-fixed in 4% PFA for 12 hours. In embryonic pulse-chase experiments, E10 – E13 embryos were lightly fixed by immersion in 4% PFA starting 8-24 hours after TM induction (as indicated in figures) and frozen for sectioning after overnight cryoprotection in 30% sucrose. Later stage embryos (E14 – E18) were perfused with 4% PFA using a 30-gauge needle and 20-cc syringe before being post-fixed in 4% PFA for 6 hours and embedded in 1.8% agarose for vibratome sectioning. Brain sections were then processed histochemically as described below.

#### Tamoxifen induction and BrdU birthdating

To reduce toxicity to pregnant females and enable sparse cellular labeling, pregnant female mice were gavaged with low-dose tamoxifen (dissolved in corn oil by gentle agitation for 24 hours), using an 18-gauge 50 mm gavage tool. Doses were in the range of 0.150 mg/30 g body weight (BW) up to 5 mg/30 g body weight (BW) (5 mg/Kg BW – 165 mg/Kg BW), with most experiments performed using doses below 1.5 mg/30 g BW. A single pulse of BrdU or EdU was intraperitoneally injected into pregnant mice at a dose of 5 mg/100 g BW, staggered before or after the tamoxifen induction depending on the goals of the experiment and driver line being used.

#### Histology and immunostaining

Fixed brains were serially sectioned at 55 um using a vibratome and collected into well plates. Sections were blocked in 10% normal goat serum and 0.1% Triton-X100 in 1x PBS, then incubated in primary antibodies (diluted in blocking solution) at 4°C overnight: RFP (rabbit polyclonal antibody, Rockland Pharmaceuticals), anti-Nkx2.1 (rabbit polyclonal, Santa Cruz Biotechnology), anti-Mash1/Ascl1 (mouse monoclonal, BD Biosciences), anti-BrdU (rat monoclonal Accurate Chemical), anti-Parvalbumin (mouse monoclonal, Sigma), anti-Somatostatin (rabbit polyclonal, Millipore), anti-TuJ1/BIII-tubulin1 (mouse monoclonal, Covance), anti-Ki67 (rabbit polyclonal, Vector), anti-ankyrin G (mouse monoclonal, Thermo Fisher). Sections were then rinsed and incubated with appropriate AlexaFluor dye-conjugated IgG secondary antibodies (1:800 goat anti-rabbit/mouse/rat, Life Technologies) and mounted in Fluoromount-G (SouthernBiotech). For BrdU immunostaining, sections were denatured in HCl (2 N) at 37°C for 45 minutes, then neutralized in sodium borohydrate buffer (0.1 M pH 8.5) twice for 10 minutes at room temperature before the normal staining protocol. EdU was visualized following a click-IT chemistry reaction with an Alexa Fluor dye-conjugated azide according to the Invitrogen EdU kit assay.

#### Imaging and serial reconstruction of whole mouse brains

Individual slides and cortical areas were imaged on a Zeiss LSM 780 confocal microscope (CSHL St. Giles Advanced Microscopy Center), including some featured 10x and 20x micrographs and all 40x or 63x micrographs. Serial whole brain imaging and volumetric whole brain imaging were completed using two different modalities and compared for consistency. The majority of whole brain image data were obtained by imaging serially-mounted immunostained frozen microtome sections with a Zeiss Axioimager microscope, which has built-in focus control, equipped with a Ludl motorized stage and a Hamamatsu Flash 4.0 camera (Paletzki and Gerfen, 2015). In this case semi-automated image acquisition was performed using Neurolucida software (MicroBrightField^TM^), before FIJI macros developed by Charles Gerfen were used to align, register, adjust, flatten, threshold, enhance contrast, and subtract background from whole brain multi-channel fluorescent image stacks (Paletzki and Gerfen, 2015). Similar imaging was completed on a Neurolucida Image Maker, Zeiss Axiomager M2, MBF Bioscience workstation in the Huang lab, supported by an equipment grant from the NIH (Award Number S10OD021759). To confirm accurate anatomical measurements and cell counts via volumetric imaging to allow tracing of individual neurites in 3 dimensions, the native fluorescent signal of some samples was imaged and analyzed using serial two-photon tomography with a 5 micrometer Z-step, as has been previously described [Ragan et al., 2012]. Image processing was completed using ImageJ/FIJI software with all alterations applied only to entire images.

### QUANTIFICATION AND STATISTICAL ANALYSIS

Manual scoring and counting of individual interneuron cell types were completed using Neurolucida 11 software either directly from slides on an Olympus Bx51 microscope, with cortical areas defined by the Mouse Brain in Stereotaxic Coordinates 4^th^ Edition atlas (Franklin and Paxinos) and cortical layers defined by the Allen Brain Atlas Coronal Adult Mouse Brain Reference Atlas. Areas for quantification were the following: medial prefrontal cortex (mPFC = anterior cingulate area (ACA), prelimbic area (PL), infralimbic area (IL) motor cortex (primary (M1), secondary (M2)) and somatosensory (barrel field and trunk subregions). Manual scoring and cell counts and 3D renderings made from digital image stack datasets were completed using Neurolucida 11, Imaris 8.0.2, or FIJI software. Unless otherwise indicated, quantification includes cell counts from the entire defined volume of a given cortical area (i.e. mPFC, MC, or SSC), while quantification of embryonic marker expression was done by using Z-stack confocal images of 6 representative coronal sections per sample through the rostral/caudal extent of the MGE. Cortical depth measurements were taken by measuring both the micrometer distance in Neurolucida from pia to cell as well as the full depth of cortex (pia to white matter) at the location of that cell. The normalized distribution (% depth) of these measurements was then plotted within ten bins of equal length from pia to white matter.

To quantify the co-localization of PV or BrdU with TdTomato^+^ cells, the ratio of PV^+^RFP^+^ or BrdU^+^RFP^+^ cells to total RFP^+^ cells within a given layer or cortical area was calculated and both the raw numbers of cells per region of interest and the normalized data and were reported. Graphical data include a minimum of three different brains of two different litters in all cases. Unpaired or paired Student’s T-tests were used to analyze data with two groups (see each figure legend for details), and one-way ANOVA and Tukey’s HSD tests were used for unbiased comparisons of multiple time points within a series. Results were displayed graphically and statistics completed using GraphPad Prism software. Histogram data showing ChC production across embryonic development by day were compared with nonlinear Gaussian curves in GraphPad Prism software, as described in the figure legends. Analysis of the fits between Gaussian curves and raw data was done with R^2^ values and D’Agostino & Pearson omnibus normality tests.

## ACKNOWLEDGEMENTS

We thank Hiroki Taniguchi and Songhai Shi for discussions and comments on the manuscript. We would also like to acknowledge Charles Gerfen and Pavel Osten for guidance on volumetric whole brain imaging techniques for collection of the data herein.

## AUTHOR CONTRIBUTIONS

Z.J.H conceived and organized the study. S.M.K. and Z.J.H. designed the experiments. S.M.K. carried out all initial single driver and intersectional fate mapping experiments, with contribution from M.M. R.R. contributed to genetic fate-mapping experiments and embryonic pulse-chase experiments. Fate mapping data were reviewed, scored and quantified by S.M.K, R.R., and M.M under the direct supervision of Z.J.H. G.K. contributed to characterization and birth dating of chandelier cells in fate mapping samples. Z.J.H and S.M.K. wrote the manuscript.

## DATA AVAILABILITY AND RESOURCE SHARING

Whole brain image datasets were submitted to the Biolucida image server for shared viewing and review (Tracked Directories>Huang Lab>Images). No other original software or data (sequences, structures, biological macromolecules, organisms, or microarray) requiring a repository were generated in this study. Raw data and materials are fully available by individual inquiry to the corresponding author.

Further information and requests for resources and reagents should be directed to and will be fulfilled by the corresponding author, Dr. Z. Josh Huang.

