## Supplemental Figures for "Multipotent radial glia sequentially deploy fate-restricted intermediate progenitors to diversify cortical interneurons"

### SUPPLEMENTAL ITEMS

#### Figure S1

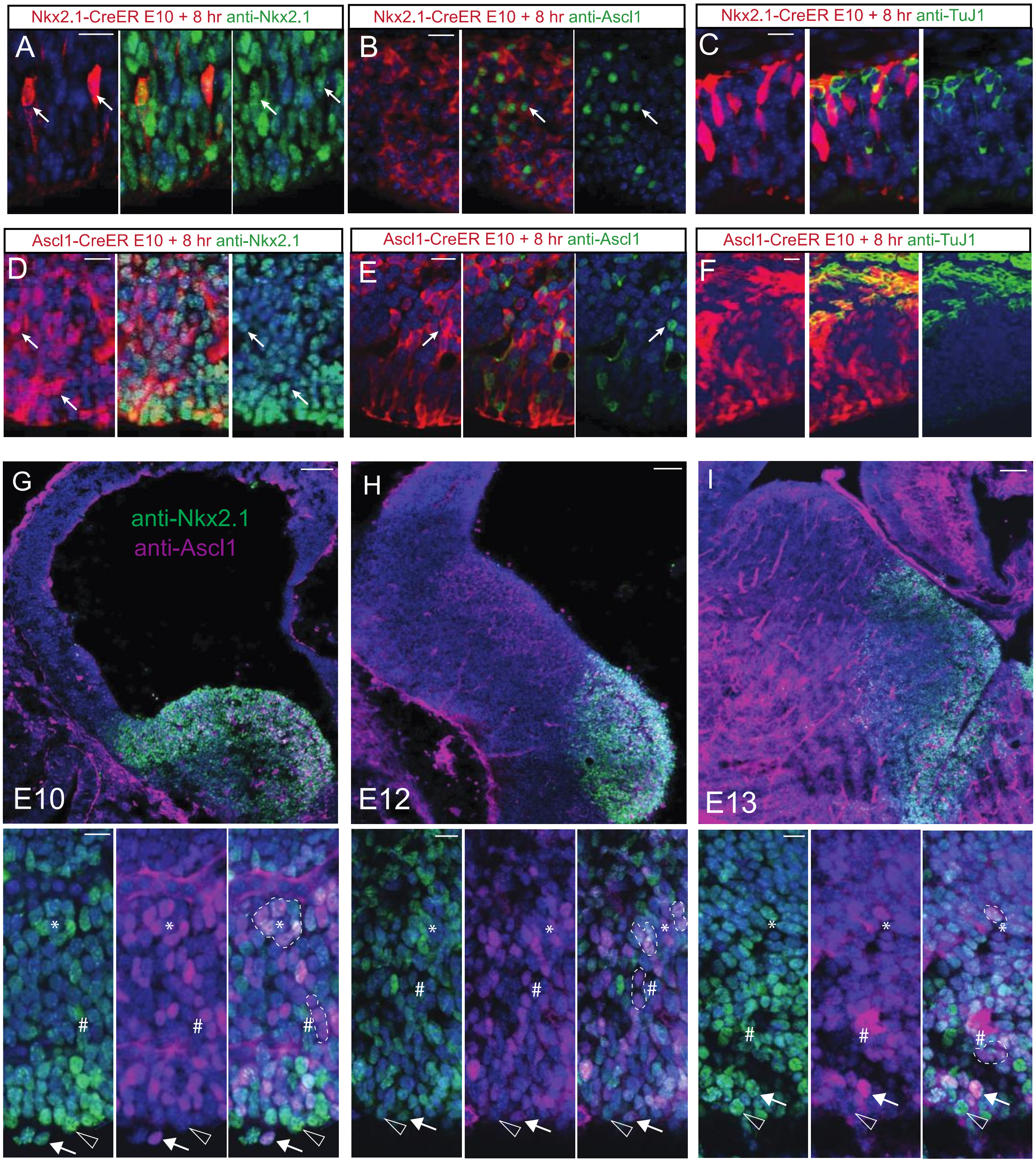

**Figure S1. Dynamic balance of Nkx2.1 and Ascl1 expression levels distinguishes multiple progenitor subtypes with different proliferative and neurogenic capacities in the MGE**

(A-C) Pulse-chase analysis of E10 Nkx2.1$^{+}$ progenitors at 8 hours showing co-labeling with anti-NKX2.1 (A, arrow) and anti-ASCL1 (B, arrow) as well as the exclusion of postmitotic marker BIII-tubulin 1 (TUJ1) (C). NKX2.1 immunoreactivity was near-ubiquitous within this VZ domain, but present at different levels in neighboring progenitors.

(D-F) Pulse-chase analysis of Ascl1$^{+}$ progenitors at 8 hours showing co-labeling with anti-NKX2.1 (D, arrow) and anti-ASCL1 (E, arrow) as well as the exclusion of TUJ1 from the cell bodies of progenitors. (F). The salt and pepper distribution of ASCL1 immunoreactivity (green) is suggestive of an oscillatory expression pattern, with *Ascl1-CreER* likely to preferentially target aIPs with sustained high-level expression.

(G-I) Distribution of anti-NKX2.1 immunopositive cells (green) in MGE at E10 (G), E12 (H), and E13 (I) demonstrates a trend for NKX2.1 expression from uniform across cell layers to be increasingly restricted to the ventricular border over time; also see Figure 1, with Ascl1 co-immunolabeled cells intermingled in the VZ and SVZ. Co-labeling of anti-NKX2.1 (green) and anti-ASCl1 (magenta) reveals a variety of progenitors with varying levels of expression of each protein. In general, high ASCL1 expression delineates an IP with limited proliferative capacity, which may be localized to the ventricular surface or deeper in the SVZ, while ventricular NKX2.1 expressing progenitors lacking ASCL1 co-expression likely represent proliferative RGs. NKX2.1 expression tends to be increasingly focused to the ventricular border over time, seen also in main Figure 1. In magnified panels at bottom, putative progenitor subtypes such as ventricular NKX2.1$^{HI}$ ASCL1$^{LO}$ RG (open arrowhead), ventricular NKX2.1$^{HI}$ASCL1$^{HI}$ aIP (arrow), non-ventricular NKX2.1$^{HI}$ASCL1$^{HI}$ aIP (*), and non-venticular NKX2.1$^{LO}$ASCL1$^{HI}$ bIP (#) are marked.

Scale bars: A-F: 10 um; G-I: 50 um in top panels, 10 um in bottom panels.

**Figure S2**

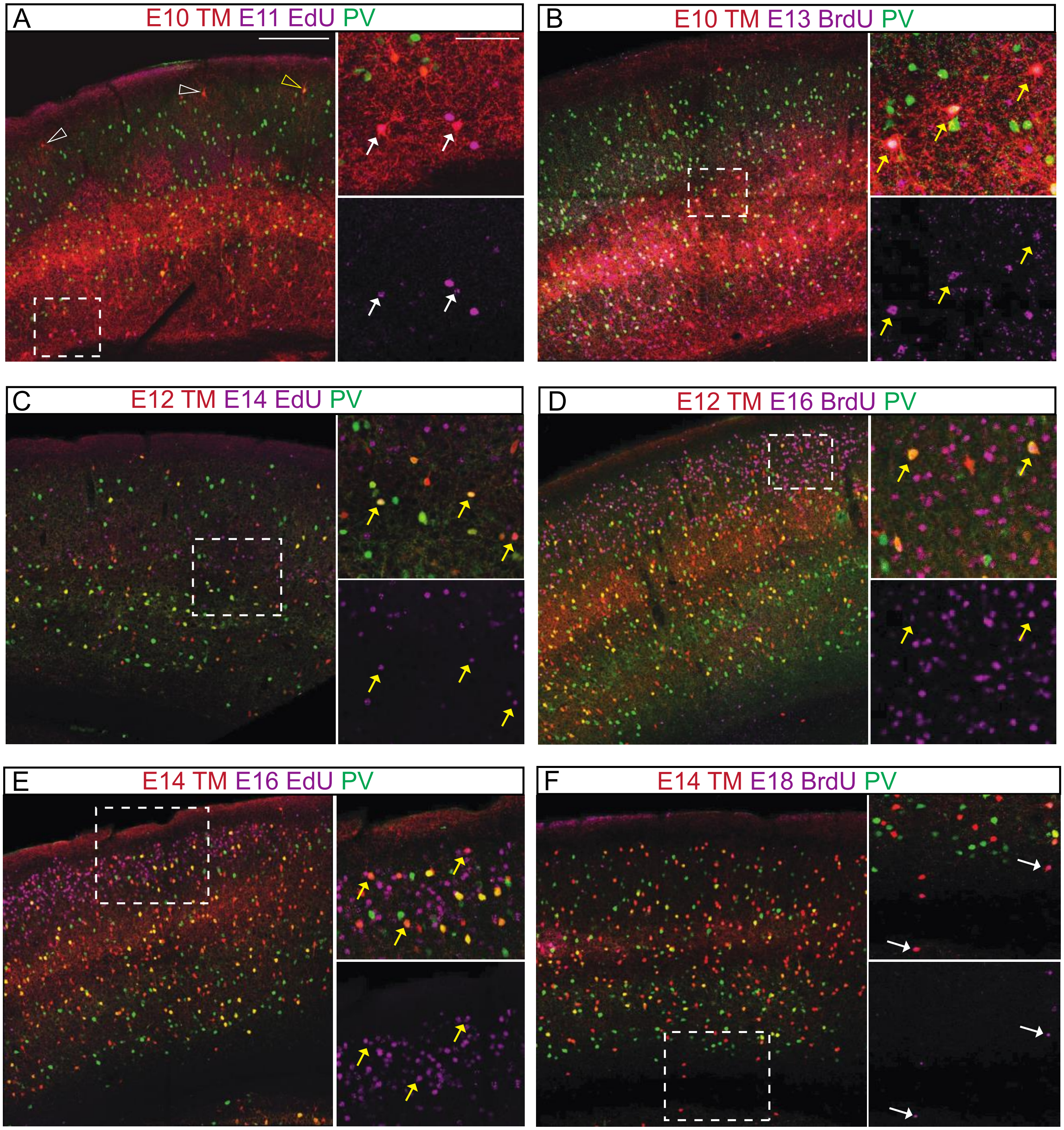
A
E10 TM E11 EdU PV
B
E10 TM E13 BrdU PV
C
E12 TM E14 EdU PV
D
E12 TM E16 BrdU PV
E
E14 TM E16 EdU PV
F
E14 TM E18 BrdU PV

**Figure S2.** ***Nkx2.1-CreER*** **fate mapping combined with cell birth dating reveal distinct waves of PV and non-PV interneuron production by layer**

(A) Early-born MGE interneurons, which were labeled by *Nkx2.1-CreER* E10 TM induction and incorporated EdU (magenta) from an E11 pulse, included primarily non-PV L5 and L6 cells (arrows in right panel, magnified from dashed box) as well as upper layer 3 ChCs (arrowheads) some were $PV^{+}$ (yellow arrowhead).

(B) Within the same cohort of cells deriving from E10 MGE progenitors, E13-born cells ($BrdU^{+}$, magenta) included a large number of L5 PV cells, indicated by yellow arrows in right panels, magnified from dashed box.

(C, D) TM induction at E12 captured a prolific burst of MGE interneuron production, with E14 EdU and E16 BrdU pulses revealing a large number of PV cells produced in inside-out order from middle to upper cortical layers (yellow arrows in subpanels).

(E, F) TM induction at E14 reveals continued interneuron production in upper cortical layers. Many L2 cells are co-labeled by an E16 EdU pulse (magenta, E), before a late burst of neurogenesis that populates cortical layers in an outside-to-inside manner, with many E18 $BrdU^{+}$ cells (magenta, F) noted in middle and lower cortical layers (yellow arrows).

Scale bars: A-F: 300 um in left panels, 100 um in right panels.

**Figure S3.**

| A | E10 | E12 | E13 | E14 | E16 | E17 |
|---|---|---|---|---|---|---|
| Nkx2.1-Flp<br>Ascl1-CreER | L5 BSK (++)<br>L5 MnC (+)<br>L6 LPC (+) | L3 ChC (+)<br>L4 BSK (++)<br>L5 BSK (++)<br>L5 MnC (+)<br>L5 DAC (++) | L3 ChC (+)<br>L3 BSK (++)<br>L3 MnC (+)<br>L3 non-MnC (+)<br>L4 DAC (+) | L2 ChC (+)<br>L2/3 BSK(++)<br>L3 MnC (+)<br>L3 non-MnC (+)<br>L4 DAC(+) | L2 ChC (+++)<br>L2 HAC (++)<br>L2 DSC (++)<br>L5/L6 ChC (+) | L2 ChC (+++)<br>L2 HAC (+)<br>L2 DSC (++)<br>L5/L6 ChC (++) |
| Nkx2.1-Flp<br>Dlx1-CreER | Dlx1 postmitotic<br>L5 BSK (++)<br>L5 MnC (+)<br>L6 LPC (+) | L4 BSK (++)<br>L5 BSK (+++)<br>L5 MnC (++)<br>L5 DAC (++) | L2/3 ChC (+)<br>L3 BSK (++)<br>L3 MnC (+)<br>L3 non-MnC (+)<br>L4 DAC (+) | L2 ChC (+)<br>L2/3 BSK(++)<br>L3 MnC (++)<br>L3 non-MnC (+)<br>L4 DAC (+) | L2 BSK (++)<br>L3 DAC (++)<br>L3 MnC (+) | L2 BSK (++)<br>L2 DAC (++)<br>L3 MnC (+) |

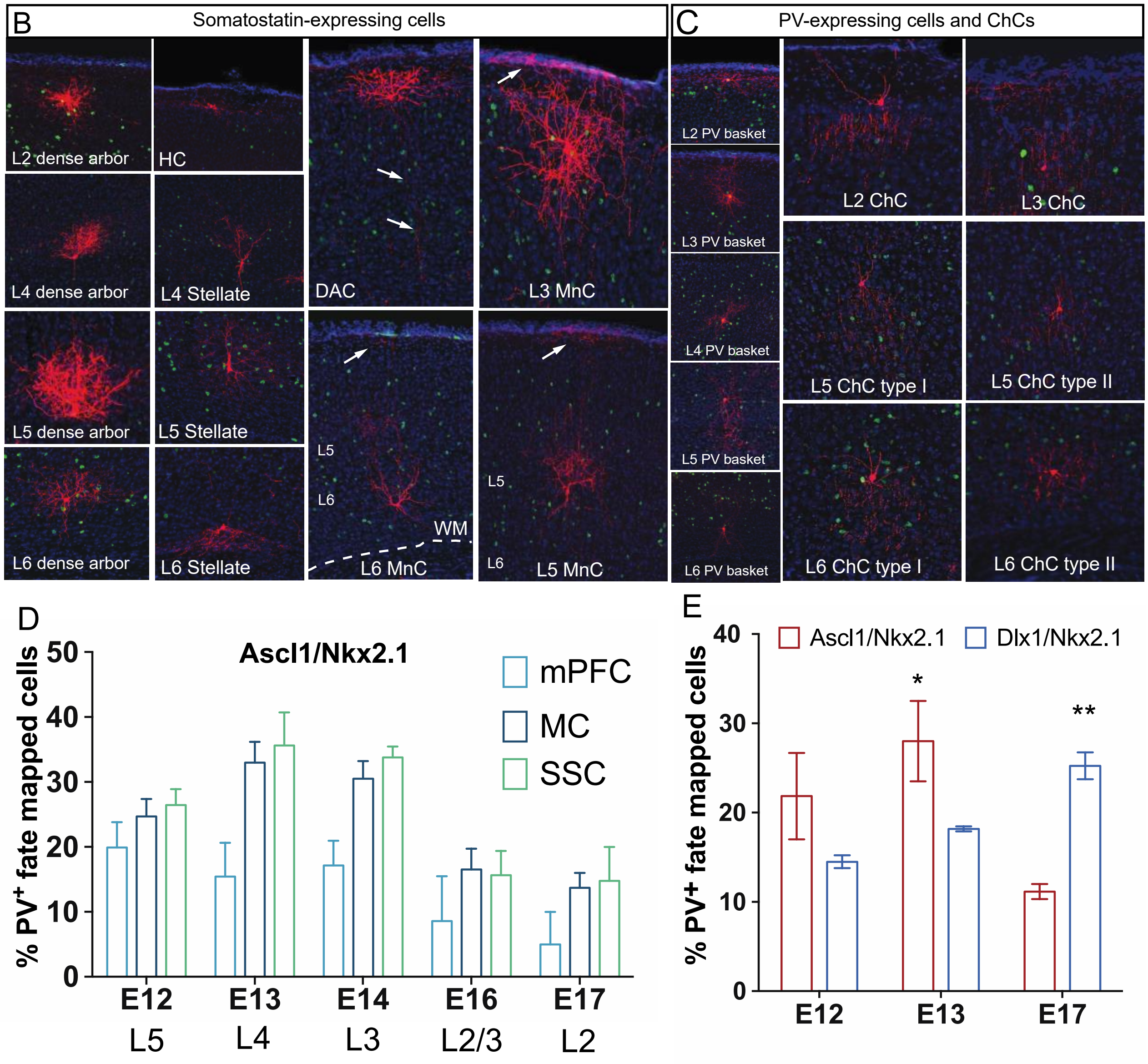

**Figure S3. IP$^{Ascl1}$ and IP$^{Dlx1}$ in MGE are sequentially allocated to specify multiple different types of interneurons**

(A) Summary of laminar and morphological interneuron types systematically generated from IPs$^{Ascl1}$ and IPs$^{Dlx1}$ across MGE neurogenesis captured by intersectional fate mapping using *Ascl1-CreER;Nkx2.1-Flp* or *Dlx1-CreER;Nkx2.1-Flp* mice. IPs$^{Ascl1}$ and IPs$^{Dlx1}$ generate cells in an inside-outside-inside sequence consisting of partially overlapping interneuron cell types during earlier embryonic times (E12-E14), but largely non-overlapping cell types at late embryonic times (E16-E18), reflecting a gradual divergence in their fate potential. BSK, basket cells; MnC, Martinottie cells, LPC, long projection cells; ChC, chandelier cells; DAC, dense arbor cells; HAC, horizontal arbor cells.

(B,C) Examples of morphological interneuron types captured by intersection of fate mapping. These are broadly divided into two non-overlapping SOM and PV-expressing populations, each includes a wide variety of laminar and morphological types such as stellate cells, DAC, HC, DSC, MnC, BSK, ChCs. These morphological types have characteristic laminar location and were captured following TM inductions at different embryonic times (L6/L5: E12 TM, L4: E13 TM, L3: E14/E15 TM, L2: E16-E18 TM; PV, green; Nissl, blue; arrows mark axons).

(D) Laminar subsets of PV$^{+}$ interneurons in mPFC, MC and SSC were produced from MGE IPs$^{Ascl1}$ on successive embryonic days between E12 – E17.

(E) IPs$^{Ascl1}$ and IPs$^{Dlx1}$ have different propensity to produce laminar subsets of PV$^{+}$ interneurons. While IPs$^{Ascl1}$ lead the generation of L4 BSKs at E13, IPs$^{Dlx1}$ dominate the production of L2/3 BSKs at E17.

**Figure S4.**

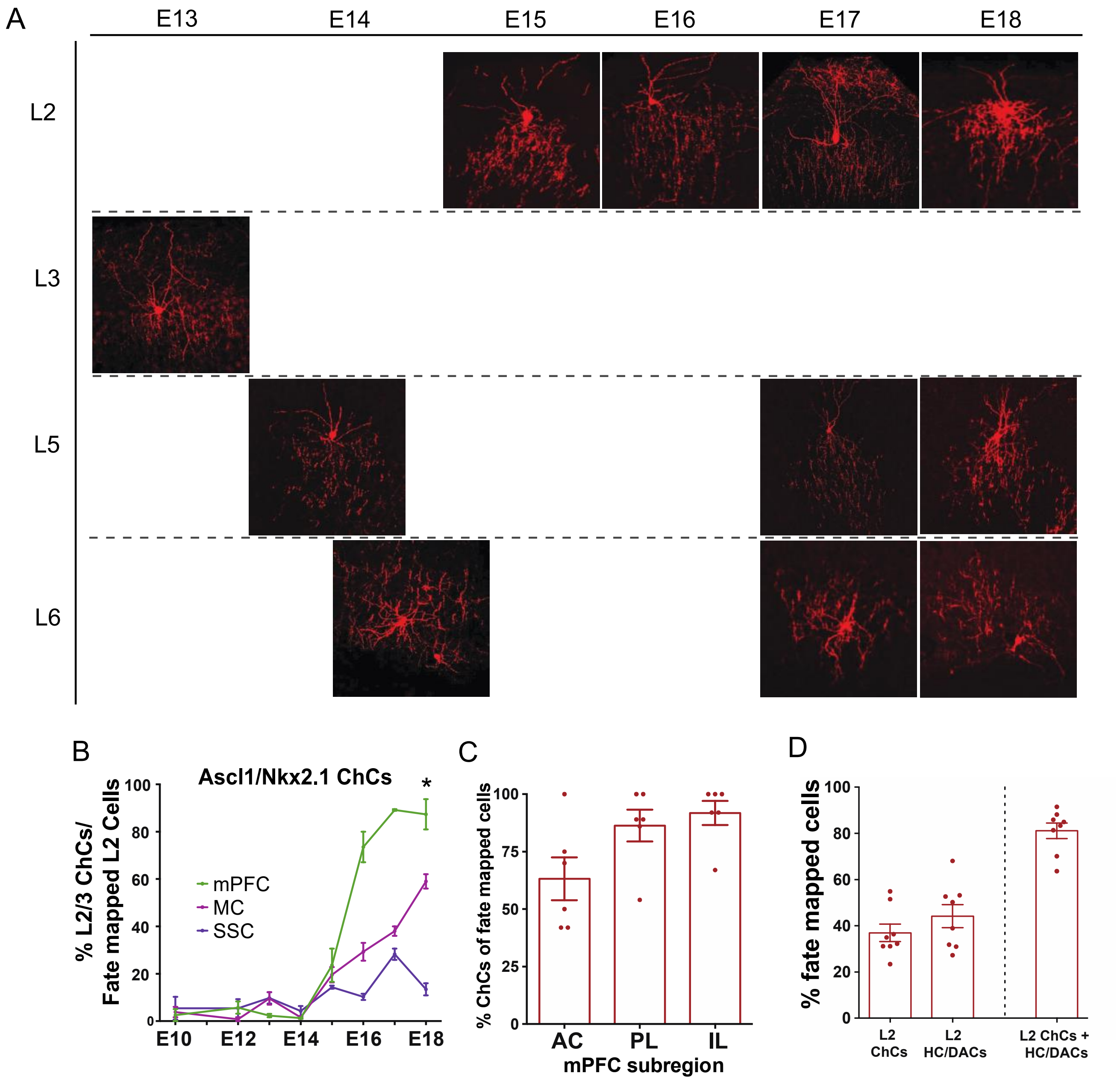
A
E13
E14
E15
E16
E17
E18
L2
L3
L5
L6
B
Ascl1/Nkx2.1 ChCs
% L2/3 ChCs/
Fate mapped L2 Cells
mPFC
MC
SSC
*
0
20
40
60
80
100
E10
E12
E14
E16
E18
C
% ChCs of fate mapped cells
0
25
50
75
100
AC
PL
IL
mPFC subregion
D
% fate mapped cells
0
20
40
60
80
100
L2
ChCs
L2
HC/DACs
L2 ChCs +
HC/DACs

**Figure S4. Two waves of ChC production are produced via a combination of direct and indirect neurogenesis**

(A) Example images the prototypical laminar ChC subtypes by birth date comprising both the early and late waves of outside-to-inside laminar settlement.

(B) IPs$^{Ascl1}$ produce a highly enriched population of ChCs in mPFC (~90% of all fate-mapped cells) relative to other cortical areas during the final stage of MGE neurogenesis (E16-18).

(C) Within mPFC, IPs$^{Ascl1}$produce a highly enriched population of ChCs in prelimbic and infralimbic cortex relative to anterior cingulate cortex during the final stage of MGE neurogenesis (E16-18).

(D) Percentage of ChCs and HC/DACs among L2 cells, accounting for ~90% of the total fate mapped cell population.
